# Cellular antibody affinity-based CRISPR Screening identifies JUNB as a broadly acting anti-viral factor

**DOI:** 10.1101/2025.06.12.659131

**Authors:** Nicole Waild, Jessica Ciesla, Xenia Schafer, Joshua Munger

## Abstract

CRISPR screening is a powerful approach to identify genetic perturbations that impact viral infection. However, most virus-focused CRISPR screens utilize selection strategies that limit the ability to identify genes important for infection. Here, we developed a novel CRISPR screening pipeline to identify cellular determinants of Human Cytomegalovirus infection based on virally induced remodeling of cellular antibody affinity (VIRCAA), which is scalable for large libraries and can identify cellular genes that impact HCMV infection at different life cycle stages. We utilized this pipeline to interrogate proteomic and transcriptomic data sets associated with the HCMV UL26 protein, which blocks anti-viral signaling during infection. We find that JUNB drives anti-viral gene expression, induces protein ISGylation, and suppresses diverse viral infections. Further, UL26 proximally interacts with JUNB and suppresses JUNB’s nuclear condensation and JUNB-mediated contraction of viral DNA replication compartments. These results highlight the VIRCAA pipeline’s utility for identifying important determinants of viral infection.

## Introduction

CRISPR screens are powerful tools used to interrogate the effects of a large number of genetic perturbations on a phenotype of interest^1,2^ and have been utilized to identify virus-host interactions that are critical for infection^3–5^. However, the success of CRISPR screening depends on the ability to segregate cellular populations based on a specific phenotype. Cellular survival has been the most frequently utilized selection in CRISPR screening, including in most virus-focused screens. However, selection for cell survival can limit the identification of cellular genes important for infection. Many genes may be important for successful infection, but their inactivation might not rescue the cellular viability of infected cells. This biases screens towards the identification of cellular genes that are important for the earliest stages of infection or towards those that inhibit cell death, potentially without impacting infection, while missing those genes that impact the later events of infection^6^. For example, the top hits of several CRISPR screens targeting Human Cytomegalovirus (HCMV) have been genes found to be important for viral entry^7–9^. To expand the space for the identification of cellular genes that impact infection, we developed a method of separating cells based on HCMV’s ability to downregulate cellular antibody affinity during infection and applied it to a targeted screen interrogating cellular genes involved in the function of the HCMV protein UL26 as a proof of concept.

HCMV, a prolific beta-herpesvirus, has a seropositivity rate of ∼50-60% in the United States, with demographic factors such as geographical location and socioeconomic status influencing this prevalence^10,11^. About 1% of newborns in the United States are infected with HCMV *in utero*, with 5-17% of these infants developing sequelae of varying severity, most notably sensorineural hearing loss and cognitive deficiencies^12^. The frequency and severity of symptoms are markedly increased in symptomatic births, with ∼90% of infants displaying one or more congenital abnormalities. Infants born to mothers who experienced primary infection during pregnancy are most at risk for symptomatic HCMV^12^. Currently, no vaccines targeting HCMV are available, and the anti-viral therapeutics used to treat HCMV exhibit poor bioavailability and toxicity, limiting their use to patients in which HCMV infection would be disabling or life-threatening^13,14^. Given these shortcomings, it is crucial to identify novel targets for therapeutic intervention by increasing our understanding of the viral and host factors that are important for infection.

HCMV is a large, enveloped virus with a ∼235kbp dsDNA genome encoding 200+ open reading frames^15^. HCMV virions are composed of a glycoprotein-studded envelope, an icosahedral capsid, and the tegument, a layer of diverse proteins located between the envelope and capsid that is unique to herpesviruses^15^. Tegument proteins are released into the host cell immediately upon infection, stimulating a pro-viral environment. The UL26 tegument protein, which possesses short 21-kDa and long 27-kDa isoforms, has been shown to be necessary for the maximal production of viral progeny, as UL26 deletion (ΔUL26) mutants grow to significantly lower titers^16,17^. During the early stages of infection, UL26 localizes to the nucleus and may play a role in the timely expression of major immediate-early (IE) genes^16,18,19^. UL26 has also been shown to be crucial for attenuating cellular anti-viral defenses as it is sufficient to block NF-κB-mediated transcriptional activation and cytokine signaling^20,21^. At later stages of the viral lifecycle, UL26 exits the nucleus and accumulates at viral assembly compartments where it plays a role in the proper phosphorylation of other tegument proteins^18^. While strides have been made to understand UL26’s role during infection, including the generation of large datasets^21,22^, the ability to functionally test the suite of genes associated with UL26 for viral replication phenotypes has been limited.

Here, we developed a CRISPR screening pipeline based on virally-induced remodeling of cellular antibody affinity (VIRCAA) to better identify host genes of interest. Utilizing this method, we identified JUNB as a broad anti-viral factor. Inactivation of JUNB, an AP-1 transcription factor subunit, enhanced the replication of multiple evolutionarily diverse viruses including HCMV, Herpes Simplex Virus, a betacoronavirus, and adenovirus. We find that the inactivation of JUNB reduces the expression of interferon-induced genes in response to infection and that UL26 proximally interacts with JUNB and is necessary to prevent JUNB nuclear condensation, which is associated with its transcriptional activation^23^. These findings provide a template for improved identification of host factors that regulate successful viral infection and identify JUNB as a critical viral restriction factor targeted by the HCMV UL26 protein.

## Results

### Development of a UL26-targeted HCMV screening pipeline based on cellular antibody affinity

HCMV has been found to downregulate surface Fc Receptor expression^24^ (Fig. 1A), raising the prospect that uninfected cells could be separated from HCMV-infected cells based on differential IgG affinity (Fig. 1B). To examine this possibility, IgG-conjugated magnetic beads were incubated with mock or HCMV-infected cells, with subsequent analysis of the IgG-bead bound and unbound fractions. The majority of both protein and DNA from uninfected cells were found to be bound to IgG-conjugated beads (Fig.1, C and D). In contrast, cells infected with WT HCMV or a ΔUL26 mutant shifted substantially to the unbound fractions, consistent with their loss of surface Fc receptor expression (Fig.1, C and D). Notably, HSV-1 infected fibroblasts exhibited a similar downregulation of IgG affinity, whereas infection by the OC43 coronavirus had no impact on IgG-bead affinity (Fig. S1). The ability to segregate infected from uninfected cells based on differential IgG affinity has the potential to be utilized as a selection in an HCMV-specific CRISPR screening pipeline.

**Figure 1.**
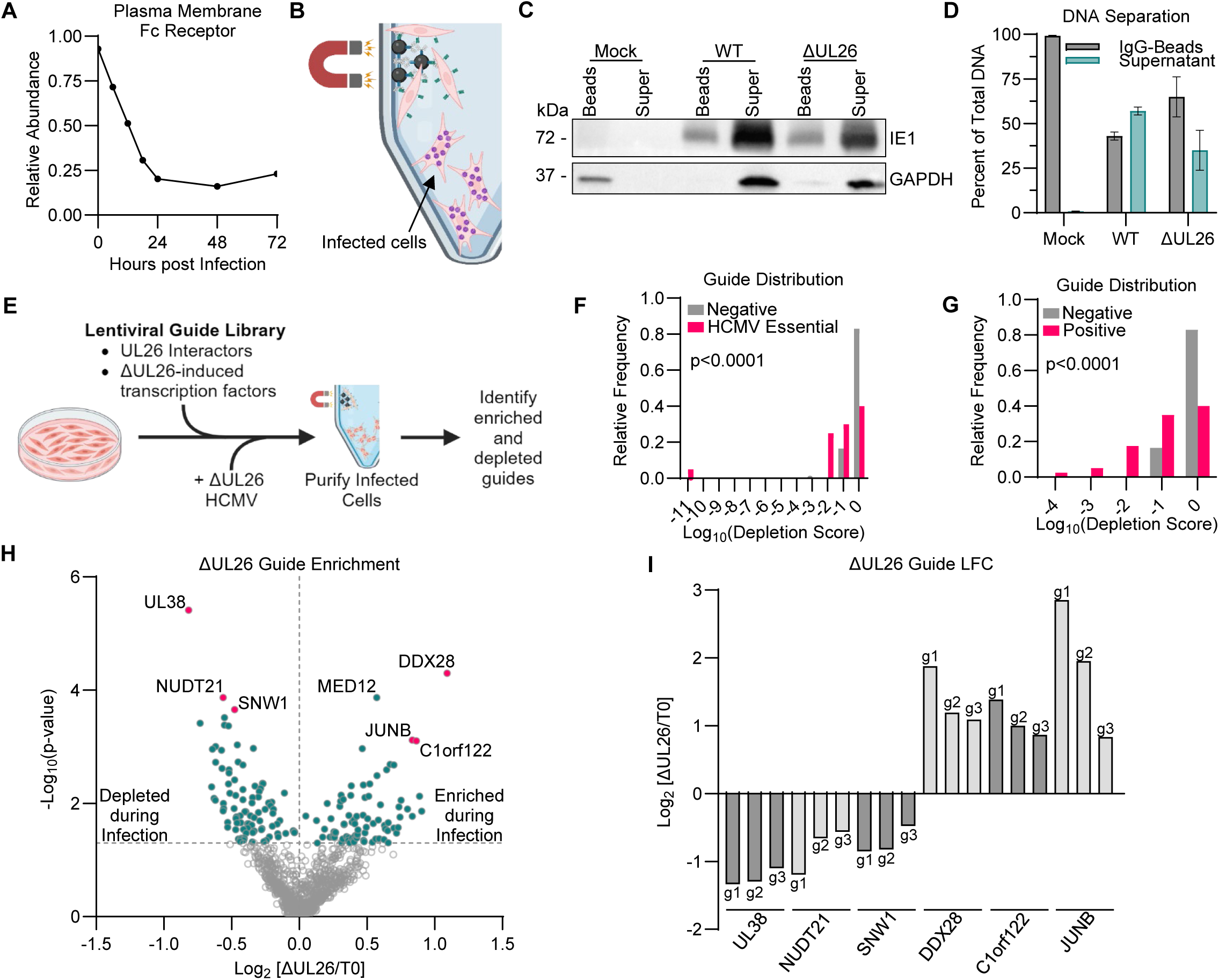
Development of the VIRCAA CRISPR screening pipeline. (A) Fc receptor expression on the plasma membrane during HCMV infection (data from Weekes et al.^23).^ (B) Schematic of cell separation based on differential Fc-receptor expression. (C,D) Mock, WT, or ΔUL26-infected MRC5 cells (MOI = 1.0) were separated based on IgG affinity at 72 hpi, and the resulting fractions analyzed for the indicated proteins (C) or DNA (D). (E) Graphical representation of the CRISPR screening pipeline. (F) Frequency distribution of the log10 depletion score of essential HCMV genes (positive) and non-targeting guides (negative). Mann-Whitney test p<0.0001. (G) Frequency distribution of the log10 depletion score of the top 40 hits from the Hein et al. screen^3^ (positive) and non-targeting guides (negative). Mann-Whitney test p<0.0001. (H) Volcano plot displaying the log_2_ fold-change (ΔUL26 vs. T0) of guides against their respective p-values. Guides that are significant (-log_10_(p-value) > 1.3) are colored teal. Guides targeting genes of interest are colored magenta and labeled. (I) log_2_ fold-change (ΔUL26 vs. T0) of the top 3 guides for the 3 most depleted and 3 most enriched genes.

The HCMV UL26 protein attenuates the expression of anti-viral gene expression^20,21^, yet the mechanisms involved remain unclear. To identify cellular genes that can modulate ΔUL26 infection, we created a lentiviral guide library based on UL26-focused datasets: a proximity-based UL26 interactome consisting of ∼600 putative UL26 interacting proteins^22^ and ∼125 transcription factors that were predicted by ChIP-X Enrichment Analysis^25^ to be activated during ΔUL26 infection based on RNA expression data of WT versus ΔUL26 infection^21^. Two sets of positive control guides were included in the library: a set of guides targeting HCMV essential genes and a set of guides consisting of the top hits from a previous HCMV-survival CRISPR screen^7^. Lastly, a set of non-targeting guides were included as negative controls. This guide library (Supplementary File 1) was transduced into MRC5 fibroblasts and subsequently mock-infected or infected with a ΔUL26 virus. After 72 hours post-infection, cells were harvested and processed for binding to IgG-conjugated beads, with the bead-bound and unbound infected cell fractions collected for deep sequencing (Fig. 1E & Supplementary File 2).

The resulting sequencing data were processed via MAGeCK RRA^26^ comparing the guide abundance in ΔUL26-infected unbound populations versus those present in guide-transduced cells prior to infection (T0) (Supplementary File 3). The majority of negative control guides were not depleted in ΔUL26-infected unbound cells, with ∼80% of non-targeting guides having a log_10_depletion score of 0 (Fig. 1, F and G). In contrast, guides targeting essential HCMV genes as well as those targeting previously described CRISPR hits^7^ were substantially depleted in ΔUL26-infected unbound cells relative to controls (Fig 1, F and G), confirming the importance of these genes to infection. These data indicate that virally-induced changes to cellular antibody affinity can segregate guides that attenuate HCMV infection from negative controls.

We next turned our attention to the UL26-focused guides that were enriched or depleted in ΔUL26-infected unbound cells. First, however, we wanted to identify any guides that might impact bead binding irrespective of infection. To this end, we identified guides that were enriched or depleted in mock-infected bead-bound populations that would be predicted to artificially impact bead binding and were therefore not analyzed further.

Of the total library population, many guides were depleted in ΔUL26-infected unbound cells, suggesting that they target pro-viral genes, whereas others were enriched in this population, suggesting that the genes they target are anti-viral (Fig. 1H). UL38, an HCMV gene that inhibits apoptosis^27^, was the most significantly depleted guide, followed by NUDT21 and SNW1, both of which act as RNA processing factors^28,29^ (Fig. 1, H and I). The most significantly enriched guides targeted DDX28, a DEAD-box helicase that enables the assembly of the mitochondrial large ribosomal subunit^30^. Additionally, the mediator complex subunit MED12^31^, the AP-1 transcription factor JUNB^32^, and C1orf122 were substantially enriched (Fig. 1, H and I). Notably, MED12 was found to impact bead binding independent of infection (Supplementary File 3) and was therefore removed from further consideration.

### JUNB restricts HCMV infection

Of the highest-scoring hits, the top two guides targeting JUNB demonstrated the greatest enrichment in ΔUL26-infected cells (Fig. 1I). JUNB forms dimers with JUN, FOS, ATF, and MAF proteins to form the transcription factor AP-1 complex^33^. While other JUN transcription factors, notably c-JUN, are considered proto-oncogenes that stimulate proliferation, JUNB is anti-proliferative, competitively inhibiting c-JUN-mediated gene activation, restricting oncogenesis, and stimulating inflammatory gene expression^32–35^.

To determine how JUNB might impact HCMV infection, we targeted its expression through the transfection of Cas9 ribonuclear proteins (RNPs) coupled to JUNB guides containing different sequences from those present in our screen. Delivery of these JUNB-specific RNPs reduced JUNB expression (Fig. 2A) and increased ΔUL26 viral spread (Fig. 2, B and C). To further validate the impact of JUNB on infection, we utilized CRISPRi to knock down JUNB transcription. CRISPRi-mediated targeting of JUNB decreased JUNB RNA abundance by ∼40% relative to controls (Fig. 2D). JUNB knockdown increased the viral spread of both ΔUL26 and WT HCMV relative to non-targeting controls (Fig. 2, E to H). Further, CRISPRi-targeting of JUNB increased the production of WT and ΔUL26 viral progeny during low MOI infection (Fig. 2I). However, at a high MOI, JUNB knockdown increased the production of WT HCMV but did not significantly impact the production of infectious ΔUL26 virus (Fig. 2J). Given that CRISPRi and CRISRPn-RNP mediated reductions in JUNB abundance were incomplete, we targeted JUNB using lentivirus and selected a monoclonal population, resulting in a greater reduction in JUNB expression (Fig. 2K and Fig. S2). Lentivirus-mediated CRISPR-inactivation of JUNB increased WT and ΔUL26 viral spread and the production of viral progeny during low MOI infection (Fig. 2L to P). Inactivation of JUNB also increased the kinetics of WT infection at a higher MOI, e.g., increasing virus production at 72hpi, although end-point titers were not substantially affected (Fig. 2Q).

**Figure 2.**
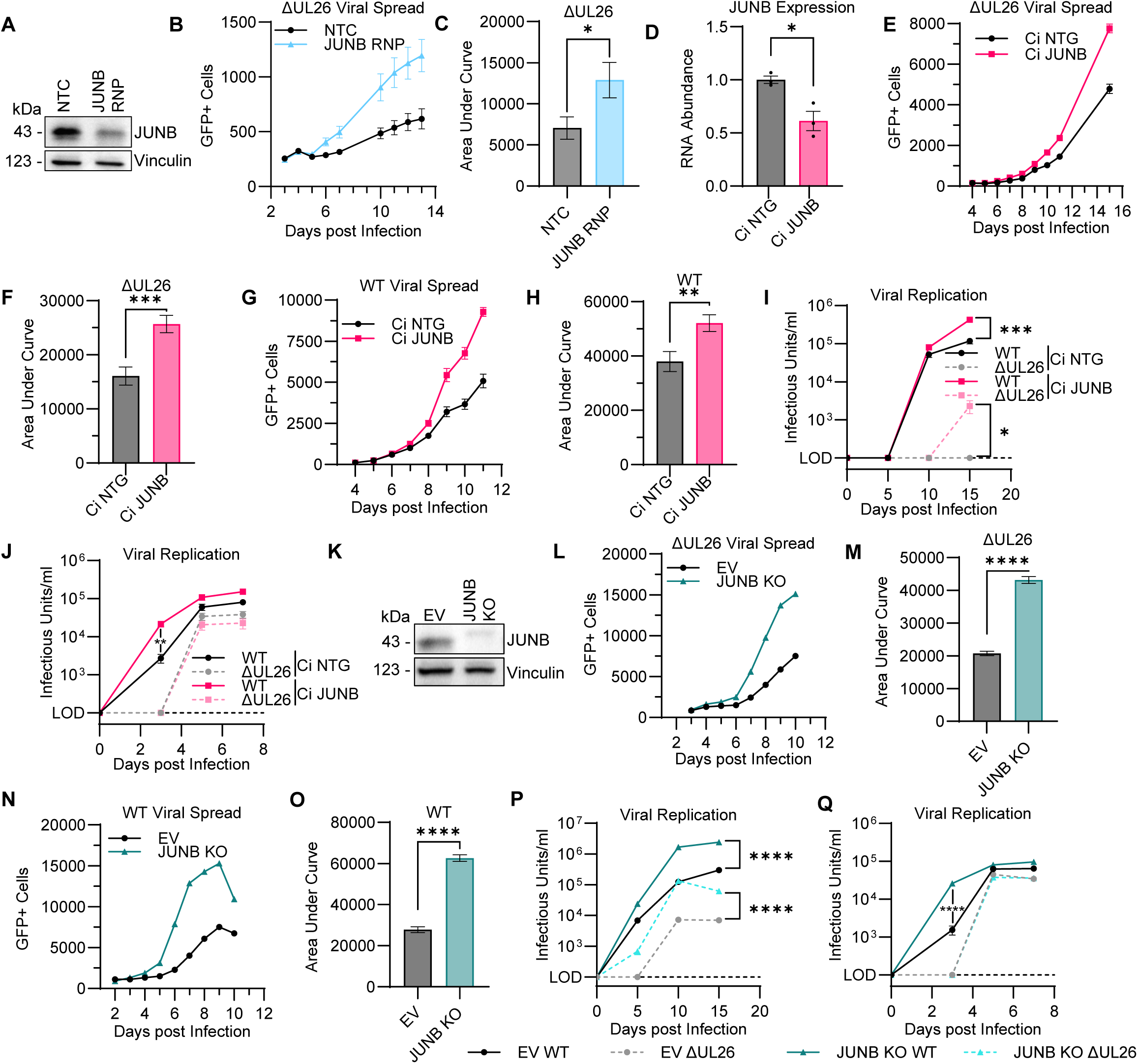
JUNB restricts HCMV infection. (A) MRC5 were transfected with CRISPR-Cas9 non-targeting control (NTC) or JUNB-targeting guide (JUNB RNP). (B-C) NTC and JUNB RNP cells were infected with ΔUL26 virus at an MOI of 0.01 and imaged for GFP expression (n = 12). (C) Area under the curve of the data presented in (B). (D) MRC5 were transduced with CRISPRi non-targeting guide (Ci NTG) or CRISPRi JUNB-targeting guide (Ci JUNB). RNA was harvested for qPCR (n = 3). (E-F) Ci NTG and Ci JUNB cells were infected as in (B) (n = 12). (F) Area under the curve of the data represented in (E). (G-H) Ci NTG and Ci JUNB cells were infected with WT virus at an MOI of 0.01 and imaged for GFP expression (n = 12). (H) Area under the curve of the data represented in (G). (I) Ci NTG and Ci JUNB cells were infected with WT or ΔUL26 virus at an MOI of 0.01, harvested at 5-, 10-, and 15-days post infection, and titered via GFP expression (n = 12). (J) Ci NTG and Ci JUNB cells were infected with WT or ΔUL26 virus at an MOI of 1, harvested at 3-, 5-, and 7-days post infection, and titered via GFP expression (n = 12). (K) MRC5 were transduced with CRISPR empty vector (EV) or CRISPR JUNB sgRNA (JUNB KO). (L-M) EV and JUNB KO cells were infected with ΔUL26 virus at an MOI of 0.1 and imaged for GFP expression (n = 24). (M) Area under the curve of the data represented in (L). (N-O) EV and JUNB KO cells were infected with WT virus at an MOI of 0.1 and imaged for GFP expression (n = 24). (O) Area under the curve of the data represented in (N). (P) EV and JUNB KO cells were infected with WT or ΔUL26 virus at an MOI of 0.1 and titered as in (I) (n = 24). (Q) EV and JUNB KO cells were infected and titered as in (J) (n = 33). ns = non-significant, * = p<0.033, ** = p<0.002, *** = p<0.0002, **** =

### JUNB overexpression attenuates ΔUL26 but not WT HCMV infection

Our results suggest that JUNB restricts HCMV infection, raising the possibility that pharmacological activation of JUNB could inhibit HCMV replication. Anisomycin has previously been found to induce the phosphorylation of JUNB^36^ and we observe that anisomycin treatment leads to transient phosphorylation of JUNB (Fig. 3A), indicative of its activation. Consistent with JUNB activation inhibiting HCMV, we find that Anisomycin treatment reduced viral replication (Fig. 3B). At significantly higher concentrations, anisomycin can inhibit translation via JNK activation^37,38^, however at the concentrations employed here, translation was not affected and minimal cytotoxicity was observed (Fig. S3). To further explore JUNB-mediated restriction of HCMV infection, we over-expressed JUNB via lentiviral transduction (Fig. 3C). JUNB overexpression reduced ΔUL26 viral spread (Fig. 3, D and E). In contrast, JUNB overexpression increased WT HCMV viral spread (Fig. 3, F and G). This trend held true for the production of infectious virus. Specifically, JUNB overexpression resulted in a slight but significant induction of WT HCMV infection, but significantly inhibited ΔUL26 infection (Fig. 3H). Collectively, these data indicate that the lack of UL26 sensitizes viral infection to JUNB overexpression; however, when UL26 is present, JUNB overexpression can slightly facilitate productive infection suggesting that UL26 modulates JUNB activity in this context.

**Figure 3.**
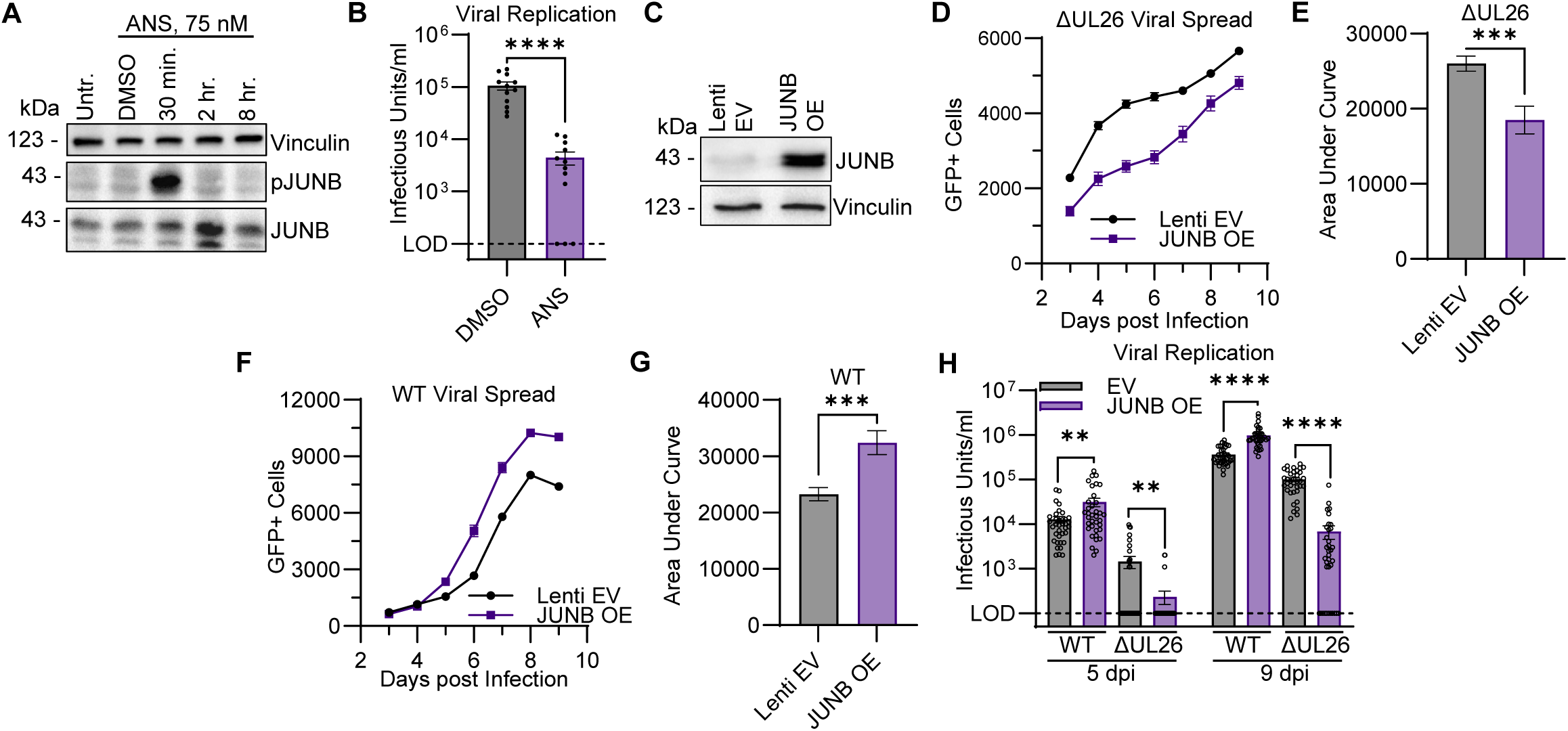
JUNB overexpression attenuates ΔUL26 viral infection. (A) MRC5 were left untreated (untr.) or treated with DMSO or 75 nM anisomycin (ANS) for the indicated time points. (B) MRC5 were infected with WT virus at an MOI of 1 and treated with 75 nM ANS at 2 hpi. Virus was harvested at 4 dpi and titered via GFP expression (n = 12). (C) MRC5 were transduced with pLenti CMV/T0 (Lenti EV) or JUNB overexpression vector (JUNB OE). (D-E) Lenti EV and JUNB OE cells were infected with ΔUL26 virus at an MOI of 0.1 and imaged for GFP expression (n = 36). (E) Area under the curve of the data represented in (D). (F-G) Lenti EV and JUNB OE cells were infected with WT virus at an MOI of 0.1 and imaged for GFP expression daily (n = 36). (G) Area under the curve of the data represented in (F). (H) Lenti EV and JUNB OE cells were infected with WT or ΔUL26 virus at an MOI of 1, harvested at 5- or 9-days post infection, and titered via GFP expression (n = 36). ** = p<0.002, *** = p<0.0002, **** = p<0.0001.

### JUNB limits HCMV viral DNA replication

JUNB was previously found in a UL26-interacting protein dataset based on proximal biotinylation^22^. To test this putative interaction, we employed a recombinant HCMV strain in which UL26 is fused to TurboID^22^. During infection with TurboID-UL26 virus, JUNB was robustly biotinylated by the UL26 fusion protein (Fig. 4A), verifying the UL26 proximity pulldown of JUNB.

**Figure 4.**
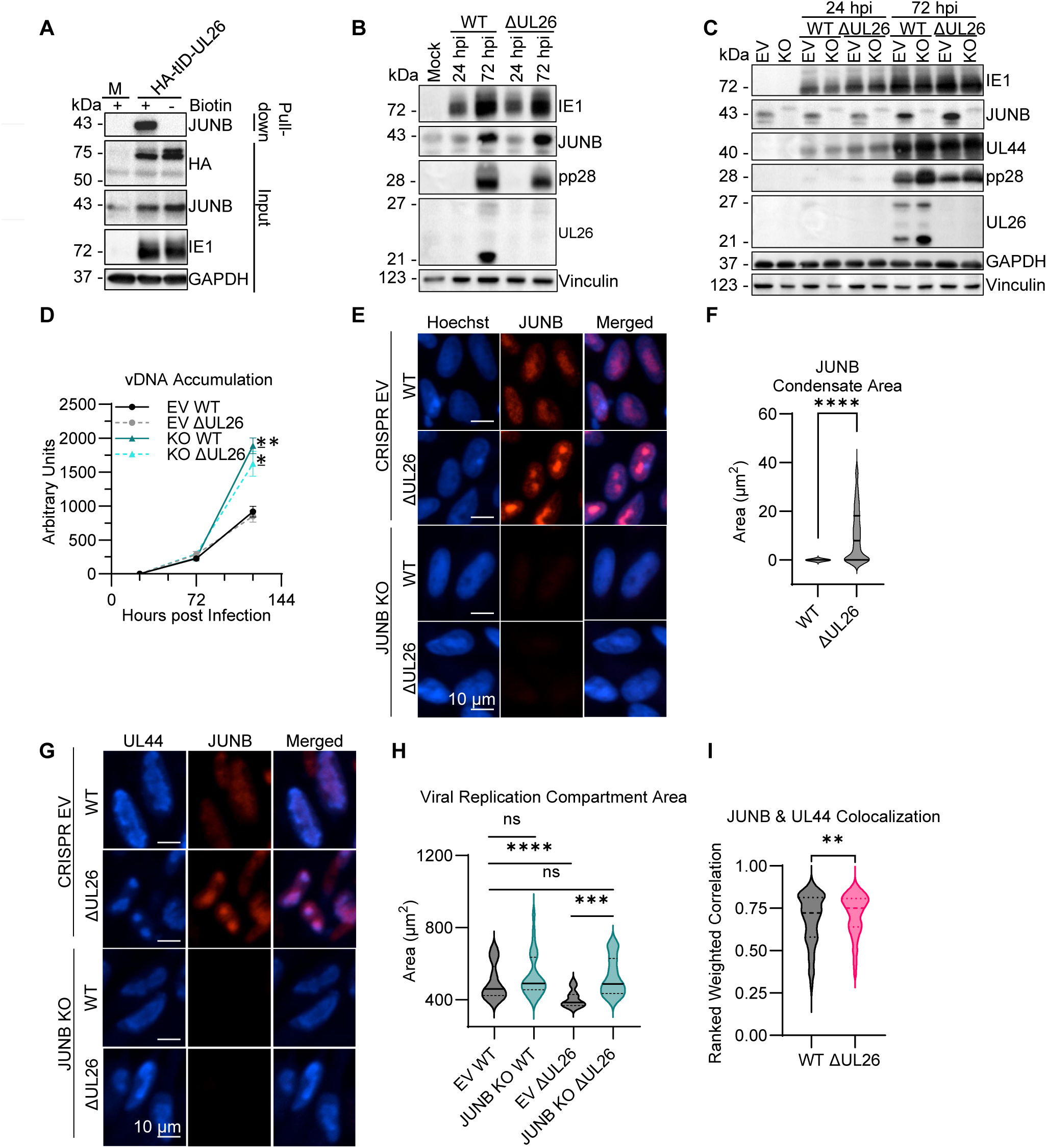
UL26 is necessary to block JUNB-mediated condensation of viral DNA replication compartments. (A) MRC5 were mock infected or infected with a HA-TurboID-UL26 (HA-tID-UL26) at an MOI of 3. (B) MRC5 were infected with WT or ΔUL26 virus at an MOI of 1. Protein was harvested 24- and 72-hpi. (C) EV and JUNB KO cells were infected as in (A) and harvested for protein at 24- and 72-hpi. (D) EV and JUNB KO cells were infected with WT or ΔUL26 virus at an MOI of 0.1 and harvested for DNA at 24-, 72-, and 120-hpi (n = 3). (E) EV and JUNB KO cells were infected as in (A). At 72 hpi, cells were imaged for nuclei and JUNB fluorescence. (F) Quantification of JUNB condensate area (n ≥ 660). (G) EV and JUNB KO cells were infected as in (A). At 72 hpi, cells were imaged for UL44 and JUNB. (H) Quantification of viral replication compartment area based on UL44 area (n ≥ 280). (I) Ranked weighted correlation of UL44 and JUNB (n ≥ 360). ns = non-significant, * = p<0.033, ** = p<0.002, *** = p<0.0002, **** = p<0.0001.

We find that JUNB accumulates during both WT and ΔUL26 infection (Fig. 4B). JUNB inactivation did not impact the accumulation of IE1, an HCMV immediate early viral gene, nor did it impact the accumulation of UL44, an early protein that functions as a viral DNA polymerase processivity factor^39–41^. In contrast, JUNB inactivation increased the accumulation of the UL26 protein, another early gene, as well as the pp28 protein (Fig. 4C), a true late gene whose levels depend on viral DNA replication. We therefore analyzed viral DNA accumulation and found that JUNB inactivation substantially increased viral DNA during both WT and ΔUL26 infection (Fig. 4D). Collectively, these data indicate that JUNB attenuates infection downstream of immediate early gene expression, reducing viral DNA synthesis. Further, these results suggest that the VIRCAA pipeline can identify cellular factors that contribute to the later stages of HCMV infection.

### UL26 is necessary to block the JUNB-mediated contraction of viral DNA compartments during infection

Upon activation, JUNB forms nuclear condensates^23^. We examined JUNB nuclear condensation during HCMV infection and found that, during WT infection, JUNB was dispersed throughout the nucleus (Fig. 4E). In contrast, in cells infected with ΔUL26, JUNB formed large nuclear condensates consistent with its activation. Upon ΔUL26 infection, the nuclear area of JUNB condensation was substantially increased relative to WT infection (Fig.4, E and F). These data indicate that UL26 prevents the formation of JUNB condensates during HCMV infection.

Given that JUNB inactivation increased viral DNA replication (Fig. 4D), we examined how the presence of JUNB impacts viral DNA replication compartments, which can be accessed via immunofluorescence of the viral DNA polymerase processivity factor UL44^39–41^. During ΔUL26 infection, viral DNA replication compartments were significantly smaller relative to those present in WT-infected cells (Fig. 4, G and H). Notably, the reduction in viral replication compartment size observed during ΔUL26 infection was reversed upon JUNB inactivation.

Further, JUNB was found to co-localize with viral DNA replication compartments during infection. This colocalization occurred regardless of whether the cells were infected with WT or the ΔUL26 mutant (Fig. 4I), although there was a slight reduction in the colocalization correlation coefficient in WT-infected cells. These data indicate that JUNB impacts the organization of viral DNA replication compartments in the absence of UL26.

### JUNB is important for anti-viral gene expression and protein ISGylation during viral infection

Infection with the ΔUL26 mutant has been found to strongly induce the expression of anti-viral cytokine signaling genes^21^ , many of which are regulated by JUNB (Fig. 5A)^42^. ISG15 expression, which is strongly induced by HCMV infection, leads to increased protein ISGylation, a post-translational modification activated by cytokine signaling and other inflammatory events^21,43^. Inactivation of JUNB substantially reduced ISG15 expression during both WT and ΔUL26 infection (Fig. 5B). JUNB inactivation also blocked HCMV-induced protein ISGylation during WT and ΔUL26 infection (Fig. 5C), suggesting that JUNB plays a critical role in the regulation of ISG15 expression and stimulation of ISG15 protein conjugation. JUNB was also found to be important for the expression of the interferon-stimulated genes (ISGs) IFIT3 and BST2, which are induced during HCMV infection^44–46^. The expression of IFIT3 and BST2 were both reduced during WT and ΔUL26 infection in JUNB KO cells relative to control cells (Fig. 5, D and E). These data indicate that JUNB is important for the induction of interferon-stimulated genes during HCMV infection and is necessary for HCMV-mediated induction of protein ISGylation.

**Figure 5.**
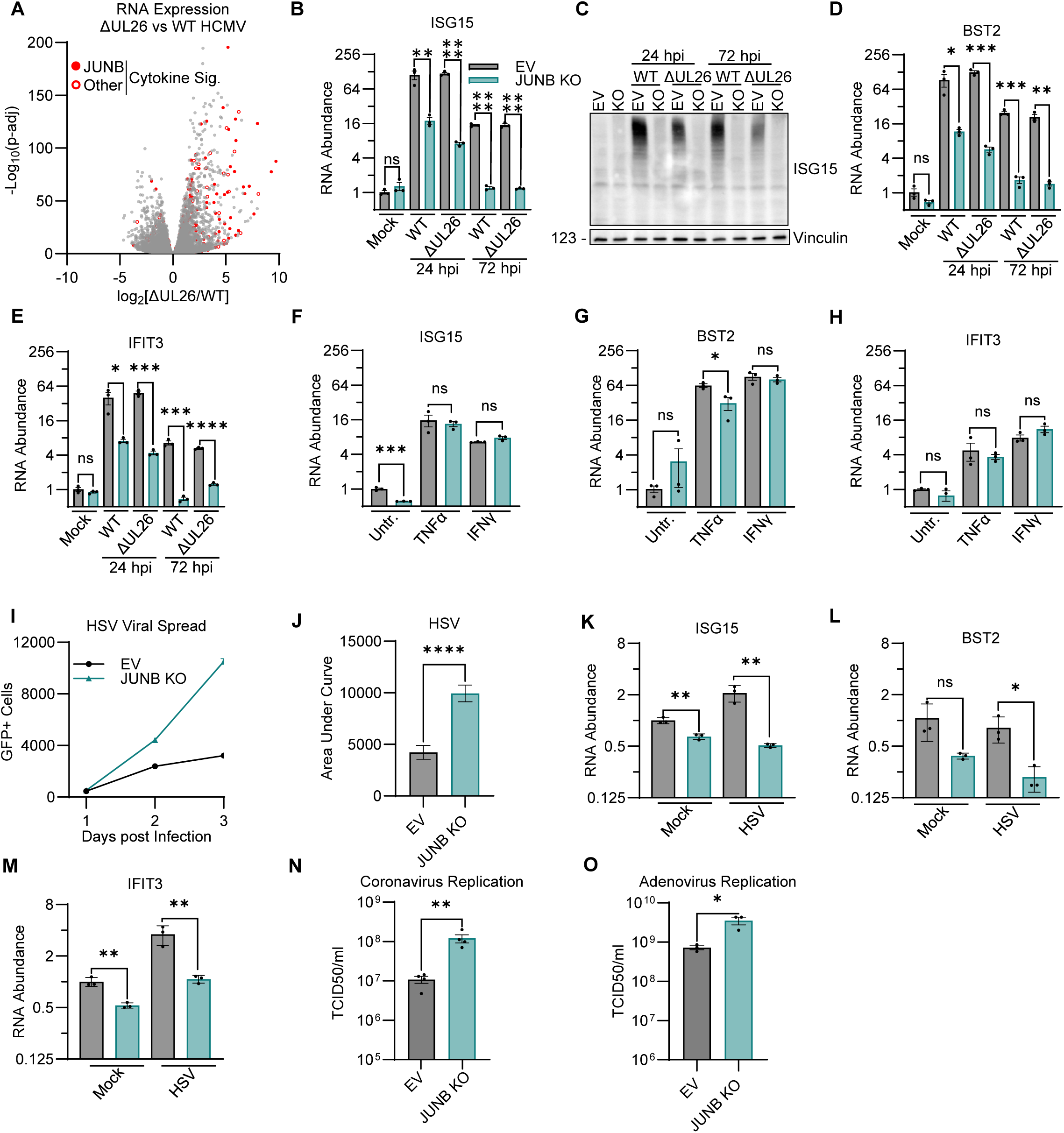
JUNB inactivation attenuates anti-viral gene expression and blocks virally induced ISGylation. (A) RNA-seq data comparing ΔUL26 vs. WT gene expression (data from Goodwin, Schafer, and Munger^21^). Cytokine signaling genes are colored red, with cytokine signaling genes that are also JUNB targets filled in red. (B-E) EV and JUNB KO cells were infected with WT or ΔUL26 virus at an MOI of 1 and harvested for protein (C) or RNA (B, D-E, n=3) at 24 and 72 hpi. (F-H) EV and JUNB KO cells were left untreated or treated with 10 ng/ml TNFα or 10 ng/ml IFNγ for 24 hours and harvested for RNA (n=3). (I-J) EV and JUNB KO cells were infected with HSV at an MOI of 0.01 and imaged for GFP expression daily (n=36). (J) Area under the curve of the data represented in (I). (K-M) EV or JUNB KO cells were infected with HSV at an MOI of 1 and harvested for RNA at 8 hpi (n=3). (N) EV or JUNB KO cells were infected with coronavirus at an MOI of 0.01. Viral supernatant was collected at 24 hpi and titered via TCID50. (O) EV or JUNB KO cells were infected with adenovirus at an MOI of 0.01, harvested for viral progeny at 11 dpi, and titered via TCID50. (M-O) Grey bars = CRISPR EV, teal bars = JUNB KO. ns = non-significant, * = p<0.033, ** = p<0.002, *** = p<0.0002, **** = p<0.0001.

### JUNB restricts the replication of evolutionarily diverse viruses

Given the findings that JUNB can regulate ISG RNA abundance during HCMV infection, we wanted to examine its role in regulating ISG expression in other contexts. We therefore evaluated how JUNB contributes to cytokine-induced ISG expression. In contrast to what was observed during HCMV infection, inactivation of JUNB had an insignificant effect on TNFα or IFNγ-induced activation of ISG15, BST2, or IFIT3 expression (Fig. 5, F to H). These results indicate that JUNB inactivation does not result in a generalized suppression of ISG expression but rather appears to be more specific for infection.

To assess whether JUNB restricts HCMV replication specifically or acts more broadly to limit viral infection, we evaluated the impact of JUNB inactivation on the replication of HSV-1, adenovirus, and the OC43 coronavirus. HSV-1 spread faster in JUNB KO cells relative to controls (Fig. 5, I and J). Additionally, consistent with what was observed during HCMV infection, the expression of ISG15, BST2, and IFIT3 was attenuated in JUNB KO cells during HSV-1 infection (Fig. 5, K to M). JUNB inactivation also stimulated the replication of OC43 and adenovirus (Fig. 5, N and O). Collectively, these data indicate that JUNB inactivation acts as a broad anti-viral restriction factor.

## Discussion

CRISPR screens are a powerful genetic tool to functionally interrogate genes for their contributions to various biological processes. Here, we developed a simple and inexpensive CRISPR-screening pipeline (VIRCAA) that can identify cellular genes important for HCMV infection and utilized it to identify JUNB as an anti-viral factor that limits the infection of numerous evolutionarily diverse viruses. With respect to HCMV infection, our data indicates that JUNB is critical for activating anti-viral gene expression and limiting viral DNA replication. Further, we find that the HCMV UL26 protein is important for limiting JUNB-mediated contraction of viral DNA replication compartments and blocking JUNB’s anti-viral activity upon overexpression.

CRISPR screening relies on the ability to isolate cellular populations based on a phenotype of interest. The majority of virally-focused CRISPR screens select for guides based on their ability to prevent virally-induced cell death. This approach can preferentially identify guide RNAs that block viral binding and entry, thereby preserving cellular viability, while not identifying guides that impact the viral life cycle at later times of infection when cells are past the point of viability^6^. For HCMV, viability-based CRISPR screening has been powerful for the identification of genes important for viral entry^7–9^. However, many clinically successful anti-viral compounds inhibit the later stages of infection. For example, ganciclovir targets viral DNA polymerase activity which inhibits infection but does not prevent the host cell’s death. Such targets would likely be missed by viability-based screening. This highlights the importance of CRISPR selection strategies that can screen for cellular determinants over a larger fraction of the viral life cycle. Our findings that JUNB does not impact the accumulation of immediate early viral genes (Fig. 4) but instead attenuates DNA replication suggests that antibody affinity-based selection is capable of identifying determinants that impact the later stages of infection and could therefore be a powerful tool in the identification of novel determinants of infection. Although our screen utilized a relatively small UL26-focused guide library, VIRCAA could be easily scaled to accommodate much larger libraries, e.g., whole genome libraries, given the ease of magnetic segregation of cells based on IgG affinity.

Virally induced remodeling of cellular antibody affinity (VIRCAA) enables the segregation of HCMV or HSV-infected cells from uninfected cells (Fig. 1 & S1). Our results demonstrate that this approach is applicable to HCMV screening, and the segregation of HSV-infected cells suggests that VIRCAA could facilitate CRISPR screening during HSV infection as well. Further, it suggests that downregulation of cell surface Fc receptors might be a shared herpesvirus phenotype. In contrast, infection with OC43, a betacoronavirus, did not impact IgG affinity (Fig. S1), and it remains to be determined whether other viral families modulate the cell surface expression of Fc receptors in ways that could be employed for CRISPR screening.

While our screen focused on UL26, our results indicate that endogenous JUNB limits both WT and ΔUL26 HCMV infection (Fig. 2). Further, JUNB inactivation attenuated interferon-stimulated gene expression during infection regardless of the presence of UL26 (Fig. 5). These results suggest that JUNB limits HCMV infection generally and plays a key role in anti-viral transcriptional responses independent of UL26. However, other JUNB-associated phenotypes only emerged in the absence of UL26. JUNB overexpression inhibited ΔUL26 replication but stimulated WT HCMV infection (Fig. 3). Further, JUNB formed condensates during ΔUL26 but not WT infection, which correlated with the JUNB-dependent contraction in viral DNA replication compartments (Fig. 4). Collectively, these data indicate that JUNB restricts HCMV infection and that UL26 modulates JUNB activity in specific contexts.

JUNB regulates gene expression through multiple mechanisms. As a transcription factor, JUNB can directly activate inflammatory gene transcription^47^, and JUNB’s transcriptional activity is shaped in part via dimer formation with different partners. JUN family members can homodimerize with other JUN family members or dimerize with various Fos^48^ and ATF^33^ proteins resulting in differential transcription programs^49^. JUNB can also impact gene expression without binding DNA through, for example, the titration and inhibition of other JUN/Fos family members^50^. Further, JUNB activity is regulated via many post-translational modifications, including phosphorylation^51,52^ and SUMOylation^53^. Notably, it has been shown that UL26 forms a complex with the E3-SUMO ligase PIAS1, which regulates inflammatory STAT signaling through STAT1 SUMOylation^54,55^, an activity that is important for limiting ISG expression during HCMV infection^56^. This, combined with UL26 proximally interacting with JUNB (Fig. 4), raise the possibility that UL26 could be regulating JUNB activity by acting as a scaffold for SUMO ligases.

During HCMV infection, JUNB localized to viral DNA replication compartments in the nucleus (Fig. 4). While this nuclear localization was diffuse during WT infection, during ΔUL26 infection both JUNB and viral DNA replication compartments substantially contracted (Fig. 4). JUNB has been reported to form nuclear liquid-liquid phase separation condensates when transcriptionally active^23^, consistent with a model in which JUNB is transcriptionally active during ΔUL26 infection. However, to our knowledge, JUNB has not been implicated in modulating viral nucleic acid replication. It remains to be determined how JUNB’s presence in viral DNA replication compartments mechanistically impacts HCMV DNA replication and whether its involvement in modulating nuclear DNA replication compartments is HCMV specific or if it impacts diverse viral infections.

We find that JUNB inactivation enhances the replication of a variety of evolutionarily diverse viruses, including cytomegalovirus, herpes simplex, coronavirus, and adenovirus (Fig. 5). JUNB’s broad activity as a viral restriction factor is consistent with our findings that JUNB is important for the induction of various anti-viral ISGs, as was observed during HCMV and HSV infection (Fig. 5). JUNB inactivation did not, however, impact ISG expression after either IFNγ or TNFα treatment (Fig. 5), indicating that JUNB’s contributions to ISG induction are not a generalized response to inflammatory stimuli, but are instead more specific to viral infection. This suggests that various molecular viral sensors could be playing a role in JUNB activation during viral infection, such as toll-like, RIG-I, or cGAS/STING signaling.

Collectively, our data indicate that VIRCAA is a powerful CRISPR pipeline to identify novel determinants of herpesvirus infection, paving the way for the exploration of new therapeutic vulnerabilities. Further, we find that JUNB is a broad-acting viral restriction factor that regulates inflammatory anti-viral responses. Understanding the mechanisms through which JUNB senses and regulates inflammatory responses could enable the modulation of these responses to limit viral infection or reduce pathogenic inflammatory response to viral infection.

## Materials and Methods

### Tissue culture and virus propagation

MRC5 fibroblasts transduced with telomerase and HEK 293T cells were cultured in Dulbeco’s modified Eagle serum (DMEM; Invitrogen #11965118) supplemented with 10% (v/v) fetal bovine serum (FBS) and 1% penicillin-streptomycin (Life Technologies #15140-122) at 37°C in a 5% (v/v) CO_2_ incubator unless otherwise stated.

AD169 viral stocks, referred to as WT HCMV in this manuscript, were the GFP-expressing BADsu-bUL21.5. The UL26-deletion virus, referred to as ΔUL26, consists of a GFP transposon insertion derived from the BAD wt clone of AD169^57^ (Genebank accession #FJ527563). Recombinant viruses expressing TurboID fused to UL26 or UL26ΔC were created as previously described^19,22^ using the BAD wt clone of AD169. All viral stocks were propagated in MRC5 fibroblasts and concentrated by ultracentrifugation. MRC5s were grown to confluence, given serum-free DMEM with 1% penicillin-streptomycin for 24 hours, and infected at an MOI of 0.01 or 0.05 for ΔUL26 stocks. Once ∼50% of the cells were detached from the culture dish, the infected cells were scraped into the media and transferred to a sterile 50 ml conical tube.

After a 5-minute 4°C centrifugation at 2,000xg, the viral supernatant was transferred to a fresh conical tube. The cell pellet was resuspended in 5 ml of viral supernatant and freeze-thawed on dry ice three times to release intracellular virus. Cellular debris was removed via centrifugation, and the supernatant was combined with the previously harvested supernatant. The viral media was mixed and distributed into ultracentrifuge tubes. Sorbitol buffer (20% D-sorbitol, 50 mM Tris-HCl, 1 mM MgCl_2_) was underlaid in each tube. Viral stocks were ultracentrifuged at 48,000xg for 1.5 hours at 23°C. The supernatant was aspirated, and the virus pellet was resuspended in a filter-sterilized solution of 50:50 3% BSA in PBS:DMEM with 10% FBS. The final viral stocks were aliquoted into screw-top vials, flash-frozen, and stored at -80°C. Stocks were titered via plaque assay.

### Viral Infections

For HCMV and HSV-YFP^58^ infections, cells were grown to confluence and given serum-free DMEM with 1% penicillin-streptomycin for 24 hours. The medium was replaced with viral adsorption media containing the indicated viral strain and MOI. After the 2-hour adsorption period, the medium was aspirated and replaced with fresh serum-free media with 1% penicillin-streptomycin unless otherwise stated.

For the coronavirus viral titer experiment, CRISPR EV and JUNB KO cells were grown and infected with OC43 virus at an MOI of 0.01 as above except media containing 2% FBS and 1% penicillin-streptomycin was used instead of serum-free media and cells were incubated at 34°C.

For the adenovirus viral titer experiment, CRISPR EV and JUNB KO cells were grown and serum-starved as above and infected with VR-5 (ATCC, VR-5) virus at an MOI of 0.01. The viral adsorption medium was removed at 1.5 hours post-infection and replaced with fresh serum-free media with 1% penicillin-streptomycin.

### Western blotting

Protein samples were harvested in disruption buffer (2% SDS, 50 mM Tris pH 7, 5% β-mercaptoethanol, 2.5% (v/v) glycerol, bromophenol blue) and detected as previously described^59^. The following antibodies were used following the manufacturer’s instructions: GAPDH (Cell Signaling, 5174S), JUNB (Cell Signaling, 3753S), HA (Santa Cruz Biotechnology, Inc., sc-7392), Vinculin (Cell Signaling, 13901S), ISG15 (Santa Cruz Biotechnology, Inc., sc-166755), phospho-JUNB (Cell Signaling, 8053S), and viral proteins IE1^60^, UL26^18^, pp28^61^, and UL44 (SeraCare, VS-P1202-2).

### CRISPR library composition, construction, and amplification

The UL26 focused CRISPR library consisted of a combination of genes predicted to interact with UL26 and genes for transcription factors predicted to be activated during ΔUL26 infection. The UL26 interacting gene set included high confidence putative UL26 interacting genes from a TurboID-UL26 proximity screen^22^ combined with UL26-interacting genes from global HCMV interactome dataset^62^. The included transcription factors were predicted to be differentially active during ΔUL26 infection based on analysis of RNA-seq data^21^. These transcription factors were identified through the analysis of transcripts that were either increased or decreased during ΔUL26 relative to wildtype HCMV infection (adj p-value<0.01 & |log2FC| >1) via ChIP-X Enrichment Analysis^25^, as implemented in ENRICHR^63^. In addition the CRISPR library contained three sets of positive controls, first, the top 40 genes from a previous HCMV-associated CRISPR screen^7^, second, 20 essential HCMV genes^57^, and third, the intersection of genes that were hits from previous CRISPR screen^7^ and high confidence cellular interactors from the HCMV interactome^62^. Each gene in the library was input to CRISPick^1,2^ using the Human GRCh38 reference genome, CRISPRko mechanism, and SpyoCas9 enzyme. Five guides were generated per gene, except for HCMV essential genes which received 10 guides per gene. CRISPOR^64^ was used to generate 200 non-targeting guides for negative controls. Duplicate genes were removed to prevent over-representation, leading to a total of 5,710 guides across all genes included.

An oligo pool of the guides with flanking sequences (5’-GGAAAGGACGAAACACCG*X*GTTTTAGAGCTAGAAATAGCAAGTTAAAATAAGGC - 3’, X being the guide sequence) was ordered from GenScript. The pooled oligo library was amplified using the primers ArrayF 5’ – TAACTTGAAAGTATTTCGATTTCTTGGCTTTATATATCTTGTGGAAAGGACGAAACA CCG – 3’ and ArrayR 5’ – ACTTTTTCAAGTTGATAACGGACTAGCCTTATTTTAACTTGCTATTTCTAGCTCTAAAAC – 3’and NEBNext high fidelity PCR master mix (New England Biolabs, M0541S). PCR reactions were pooled, run on a 0.8% agarose gel, and gel extracted using the Monarch gel extraction kit (New England Biolabs, T1020S).

The plasmid plentiCRISPRv2 (Addgene, #52961) was used as the vector for the CRISPR screen. The vector was digested with Esp3I (Thermo Scientific, ER0452) at 37°C for 1 hour. After the reaction was complete, digests were pooled and run on a 0.8% agarose gel overnight and extracted using the Monarch gel extraction kit.

Gibson assembly was used to ligate guides into the digested vector. A ratio of 330 ng of vector to 50 ng of pooled, amplified guides were used with Gibson assembly master mix (New England Biolabs, E2611L). The reactions were incubated at 50°C for 1 hour. Next, the reactions were pooled and isopropanol precipitated. The Gibson assembly reactions were mixed with isopropanol, GlycoBlue Coprecipitant (Invitrogen, AM9515), and 5 M NaCl, vortexed, and incubated at room temperature for 15 minutes. Samples were centrifuged for 15 minutes at >15,000 g at room temperature. The supernatant was aspirated, and the pellet was washed twice with ice-cold 80% (v/v) ethanol. After air-drying to remove residual ethanol, samples were resuspended in TE buffer and stored at -20°C.

The pooled library was electroporated at 100 ng/µl using Endura ElectroCompetent cells (Lucigen, 60242-1) according to the manufacturer’s instructions. After the 1-hour recovery period, cells were pooled and plated onto prewarmed LB Lennox plates with 150 ug/ml ampicillin. Plates were incubated at 37°C for 14 hours. Colonies were harvested and prepped using the NucleoBond Xtra Maxi kit (Macherey-Nagal, 740414.10) according to the manufacturer’s instructions. The library was quantified and stored at -20°C.

### Transfections

HEK 293T cells were grown to 50-70% confluence in tissue culture dishes. For each plate, PBS was mixed with pMD2.G (Addgene, #12259), psPAX2 (Addgene, #12260), and the lentiviral plasmid of interest. After a 5-minute incubation, FuGENE (Promega, E2691) was added to the PBS/DNA mixture and incubated at room temperature for 20 minutes. This solution was added dropwise to each plate of HEK 293T cells, and plates were incubated at 37°C overnight. The next day, the transfection medium was aspirated and replaced with fresh DMEM supplemented with 10% FBS and 1% penicillin-streptomycin. Plates were allowed to incubate overnight. Finally, lentiviral media were aspirated with a sterile syringe, filtered through a 45 µm syringe filter, aliquoted into screw-top vials, flash-frozen, and stored at -80°C.

### Lentiviral Titering

To titer the CRISPR library lentivirus, MRC5 fibroblasts were seeded at 2×10^5^ cells per well into 6-well plates and incubated at 37°C overnight. Three wells were harvested and counted using trypan blue (VWR, K940-100ML) and a Bio-Rad TC10 automated cell counter to calculate the doubling time of MRC5s. The remaining wells were aspirated and replaced with 0.6 ml, 0.8 ml, 0.9 ml, 0.95 ml, 0.975 ml, or 1 ml of media with 8 ug/ml polybrene (MilliporeSigma, TR1003G). To each respective well, 400 µl, 200 µl, 100 µl, 50 µl, 25 µl, or 0 µl of lentivirus was added to bring the total volume to 1 ml. The virus was allowed to adsorb for 4 hours, then was aspirated and replaced with fresh media. Plates were incubated overnight. Next, all wells were lifted with trypsin and seeded at 10,000 cells per well in a 96-well plate, using only the inner wells. Cells were allowed to rest for 48 hours. The medium was aspirated, and half of the wells received normal media and the other half media containing 1 ug/ml puromycin (MP Biomedicals, 0210055210). Cells were allowed to incubate until all cells in the -virus / +puromycin wells were dead.

The cells were washed with PBS and stained with 1x Hoechst 33342 (Invitrogen, H3570). After a 15-minute incubation, nuclei were counted using nuclei count using a Cytation 5 imaging reader (BioTek) with a 2.5X magnification lens. Fluorescence was measured using a DAPI channel with an excitation wavelength of 377 nm and an emission wavelength of 447 nm. Each nucleus was masked and counted using the Gen5 3.11 (BioTek) software. The nuclei count was used as a proxy for cell count per well. The average of replicate wells for each virus/puromycin condition was calculated and used to determine the viral titer for each dilution using the formula: [cell count after 24 hours based on doubling time * (transduced cells/total cells)] / viral supernatant volume. The average titer across all dilutions was calculated and reported as infectious units per microliter.

### Transductions

Generally, MRC5 fibroblasts were grown to 50-70% confluence in 6-well plates. Medium was aspirated and replaced with viral inoculum containing lentivirus, polybrene, and media. Plates were incubated at 37°C overnight. The viral inoculum was removed and replaced with fresh growth media, and cells were allowed to rest overnight. At 48 hours post-transduction, 1 ug/ml puromycin was added to the transduced cells. Once all cells in an equivalent, non-transduced well were dead, the transduced cells were expanded into a 10 cm dish.

For the CRISPR screen, 2×10^6^ MRC5s were seeded into 15 cm dishes. The following day, medium was aspirated and replaced with enough library lentivirus to achieve an MOI of 0.3. The total volume was brought to 10 ml, and 8 ug/ml polybrene was added. Cells were incubated at 37°C for 4 hours, then the viral inoculum was aspirated and replaced with fresh growth media. Cells were allowed to rest for 48 hours. Medium was aspirated and replaced with media containing 1 ug/ml puromycin, and cells were incubated until an equivalent, non-transduced 15 cm dish of cells died completely. Transduced cells were then split 1:2 into new 15 cm dishes.

### Magnetic bead preparation

Dynabeads M-450 Epoxy (Invitrogen, 14011) were conjugated with mouse IgG (Rockland Immunochemicals, 010-0102-0005) according to the manufacturer’s instructions. Briefly, beads were resuspended and transferred to a labeled tube, where they were washed with buffer 2 (Ca^2+^ and Mg^2+^ free PBS supplemented with 0.1% BSA and 2 mM EDTA, pH 7.4) using a magnetic strip stand. The tube of beads was then placed on a magnetic stand and the supernatant was discarded. The beads were resuspended in 980 µl of buffer 1 (0.1 M sodium phosphate buffer, pH 7.6) with an additional 20 µl of IgG (200 µg). The mixture was incubated at room temperature for 15 minutes, then BSA was added to 0.1% (w/v). The tubes were placed on a rotator and incubated at room temperature overnight. The following day, the tubes were placed on a magnetic stand, and the supernatant was discarded. The beads were washed three times using buffer 2. Once washed, the beads were resuspended in enough buffer 2 to achieve 4×10^8^ beads/ml. Coupled beads were stored at 4°C.

### Magnetic bead cell separation

Library-transduced and puromycin-selected MRC5s were grown to 90% confluence and given serum-free media for 24 hours. The plates were then infected at an MOI of 1 with ΔUL26 virus in serum-free media or left uninfected to function as mock samples. After the 2-hour adsorption period, the viral inoculum was removed and replaced with fresh serum-free media and incubated at 37°C. Some of the uninfected plates were harvested on the same day as the infection to function as t0 samples and were not processed using the magnetic beads. At 72 hours post-infection, all remaining plates were processed using the IgG-conjugated magnetic beads as described below.

IgG-conjugated magnetic beads, 50 µl per 15 cm dish, were transferred to a fresh tube and washed using buffer 2. After washing, the beads were resuspended in a 1:1 volume of buffer 2 and set aside at 4°C. To prepare the cell samples, cells were trypsinized with TrypLE (Gibco, 12605010) for 2 minutes at 37°C. The trypsin was inactivated with serum-containing media, and the cell solution was transferred to a sterile conical tube. The samples were centrifuged for 5 minutes at 500xg and 4°C. The supernatant was aspirated, and the cell pellet was resuspended in buffer 2. Samples were centrifuged for 5 minutes at 500xg and 4°C again. The buffer 2 wash was aspirated, and the cell pellets were resuspended in 1 ml of buffer 2. The cell mixture was transferred to a sterile 2 ml tube and placed on ice.

To separate the cells using the magnetic beads, 50 µl of IgG-conjugated, washed magnetic beads were added to each 1 ml sample. An additional 1 ml of buffer 2 was added to each tube, for a total volume of ∼2 ml. The samples were mixed gently by inversion and placed on a rotator at 4°C for 25 minutes. After incubation, the tubes were placed on a magnetic stand. The supernatant containing the unbound cells was transferred to a fresh 2 ml tube and centrifuged for 3 minutes at 5,000xg. The supernatant from the spun tubes was removed, and the cell pellets were resuspended in 1 ml of QuickExtract DNA extraction solution (Lucigen, QE09050) and processed using the manufacturer’s instructions. The magnetic beads containing the bound cells were gently washed three times using 1 ml of buffer 2. The wash supernatants were discarded. The bead-bound cells were resuspended in 1 ml of QuickExtract DNA extraction solution and processed using the manufacturer’s instructions. DNA samples were stored at - 80°C.

### Next-generation sequencing and processing

To check the guide representation in the plasmid library prior to transfection, the plasmid was PCR amplified using the primers NGS F1 (5’-AATGGACTATCATATGCTTACCGTAACTTGAAAGTATTTCG – 3’) and NGS R1v2 (5’ – TCTACTATTCTTTCCCCTGCACTGTTGTGGGCGATGTGCGCTCTG – 3’) and Herculase II polymerase (Agilent, 600675). The resulting product was used as the template for a second round of PCR using the primers NGS R2 (5’ – CAAGCAGAAGACGGCATACGAGATGTGACTGGAGTTCAGACGTGTGCTCTTCCGATCTTCTACTATTCTTTCCCCTGCACTGT – 3’) and NGS F2 001 (5’ – AATGATACGGCGACCACCGAGATCTACACTCTTTCCCTACACGACGCTCTTCCGATCTTAGCGCTAGTCTTGTGGAAAGGACGAAACACCG -3’) and Herculase II polymerase. The guide library was run on a 2% agarose gel and extracted using the Monarch gel extraction kit.

The amplified libraries were hybridized to the Illumina flow cell and sequenced using the MiSeq sequencer (Illumina). Single-end reads of 75nt were generated for each sample to obtain approximately 500x coverage per guide. Sequenced reads were de-multiplexed with bcl2fastq (2.19.1), and adapter trimmed to the 20bp barcode sequence using cutadapt (v1.18) before mapping to the reference sequences with Bowtie (v1.2.2)^65,66^. FeatureCounts from the subread package (v1.6.1) was then used to derive counts for each sgRNA barcode separately and to aggregate counts for each gene^67^.

To sequence CRISPR screen samples, the DNA samples were thawed and PCR amplified using the primers NGS F1 and NGS R1v2, and Herculase II polymerase. The PCR product was then used as the template for a second round of PCR using the primers NGS R2 and a unique forward primer for each sample NGS F2 (5’ – AATGATACGGCGACCACCGAGATCTACACTCTTTCCCTACACGACGCTCTTCCGATCT(1-9bp variable length sequence)(8bp barcode)TCTTGTGGAAAGGACGAAACACCG – 3’) and Herculase II polymerase. The product was run on a 2% agarose gel, and the library band was gel extracted using the Monarch gel extraction kit.

The amplified libraries were sequenced and analyzed as described above using the NextSeq550 sequencer (Illumina). Guide scores were computed using MAGeCK RRA^26^ and further analyzed using R. Briefly, the gene-wise guide scores comparing UL26 unbound to t0 and mock bead-bound to t0 were extracted into R and filtered for both negative and positive scores less than 0.02. Any genes that were present in both the mock bead-bound enriched filtered dataset and the UL26 unbound depleted filtered dataset, and vis versa, were tagged for removal. The remaining gene lists were combined into one and noted whether they were enriched or depleted.

### CRISPR-Cas9 ribonucleoprotein transfections

Gene knockout using CRISPR-Cas9 ribonucleoprotein (RNP) was performed with the Neon Transfection System 10 µl kit (Invitrogen, MPK1025) as previously described^68^. The electroporation settings used were: 1,100 V, 30 ms width, 1 pulse. The guide RNAs were ordered from Synthego and are as follows: negative control scrambled sgRNA (modified) #1 (5’ - GCACUACCAGAGCUAACUCA – 3’), JUNB (Gene Knockout Kit v2) sgRNA1 (5’ – CGCCCGGAUGUGCACUAAAA – 3’), JUNB sgRNA2 (5’ – ACGGGAUACGGCCGGGCCCC -3’), and JUNB sgRNA3 (5’ – GGUCGGCCAGGUUGACCGCC – 3’).

### Plasmid cloning

To generate a CRISPRi JUNB construct, guide RNA targeting JUNB^69^ was ordered with flanking sequences (Ci JUNB, 5’ – GGAAAGGACGAAACACCGCGGCTGGGACCTTGAGAGGTTTTAGAGCTAGAAATAGC–3’) to the CRISPRi lentivirus (LV) PURO plasmid (pLV hU6-sgRNA hUbC-dCas9-KRAB-T2a-puro with stuffer), gifted by the Harris Lab at the University of Rochester Department of Biomedical Genetics. The oligos were amplified using the primers Ci F (5’ – CTTGAAAGTATTTCGATTTCTTGGCTTTATATATCTTGTGGAAAGGACGAAACACCG–3’) and Ci R (5’ – CTTTTTCAAGTTGATAACGGACTAGCCTTATTTTAACTTGCTATTTCTAGCTCTAAAAC – 3’) and Herculase II polymerase. The plasmid was digested using Esp3I at 37°C overnight. The digested plasmid was run on a 1% agarose gel and extracted using the Monarch gel extraction kit. The amplified guide oligo was ligated into the linearized plasmid using Gibson assembly as described above using a 1:4 vector:insert molar ratio. The Gibson assembly product was transformed into heat-shock competent STBL3, and plasmids isolated from discrete colonies were Sanger sequenced to verify the presence of the JUNB sgRNA sequence. Lentivirus was generated and used to transduce MRC5 cells as described above.

To generate a CRISPRn JUNB construct, a guide targeting JUNB from the initial CRISPR screen was ordered individually with flanking sequences for plentiCRISPRv2 (JUNB sg486, 5’-GGAAAGGACGAAACACCGCTGAGGTTGGTGTAAACGGGGTTTTAGAGCTAGAAATAGC – 3’). The oligos were amplified using the primers ArrayF and ArrayR, and Herculase II polymerase. The plentiCRISPRv2 plasmid was digested using Esp3I at 37°C for 1 hour. After the reaction was complete, the digest was run on a 0.8% agarose gel overnight and extracted using the Monarch gel extraction kit. The amplified guide oligo was ligated into the linearized backbone using Gibson assembly as described above using a 1:4 vector:insert molar ratio. The product was transformed into heat-shock competent STBL3, and isolated plasmids were Sanger sequenced to verify the presence of the JUNB sgRNA sequence. Lentivirus was generated and used to transduce MRC5 cells as described above.

To generate a JUNB overexpression construct, JUNB was amplified out of pCS2 Flag-JunB (Addgene, #29687) using the primers plenti JUNB F (5’ – GGAACCAATTCAGTCGACTGGATCCGCCACCATGGACTACAAGGACGACGATGAC – 3’) and plenti JUNB R (5’ – CCACTTTGTACAAGAAAGCTGGGTCTAGATCAGAAGGCGTGTCCCTTGACCCCAAGCAG - 3’) and Herculase II polymerase. The plasmid pLenti CMV/T0 (Addgene, #22262) was digested using BamHI (New England Biolabs, R0136) and XbaI (New England Biolabs, R0145) according to the manufacturer’s instructions. The digest was run on a 1% agarose gel and extracted using the Monarch gel extraction kit. The amplified PCR product was ligated into the linearized pLenti plasmid using Gibson assembly as described above using a 1:8 vector:insert molar ratio. The product was transformed into heat-shock competent STBL3, and the isolated plasmids were Sanger sequenced to verify the presence of the JUNB insert. Lentivirus was generated and used to transduce MRC5 cells as described above.

### Monoclonal selection

MRC5 fibroblasts were transduced with lentivirus containing the CRISPRn JUNB sgRNA 486 construct as described above. Once confluent in a 10 cm dish, the puromycin-selected cells were serially diluted into two 96-well plates such that a portion of the wells would contain a single cell. After 24 hours, the medium was aspirated and replaced with fresh media containing 1 ug/ml puromycin. Each well was monitored, and wells that contained more than a single cell were discarded. Once confluent in the 96-well plate, clones were expanded stepwise into tissue culture plates of increasing size. When the clones were confluent in a 10 cm dish, the cells were split into a 15 cm dish as well as two 10 cm dishes to harvest for gDNA and protein. Protein was harvested and blotted for JUNB as described above. Genomic DNA was harvested using QuickExtract DNA extraction solution and processed as described above. DNA from WT MRC5 as well as from the clones was amplified using the primers F – JUNB Seq Set 1 (5’ – TACCGGAGTCTCAAAGCGCC – 3’) and R – JUNB Seq Set 1 (5’ – ACGTTCAGAAGGCGTGTCCCTT – 3’). These products were used as the template for another round of PCR using the primers F – JUNB Seq Set 2 (5’ – CGTCTCTCAAGCTCGCCTCT – 3’) and R – JUNB Seq Set 2 (5’ - GACCTTCTGTTTGAGCTGGGCC – 3’). The products from the second round of PCR were Sanger sequenced using the primers F – JUNB Seq Set 3 (5’ - AACAGCAACGGCGTGATCAC – 3’) and R – JUNB Seq Set 3 (5’ – CGGCCTTGAGCGTCTTCACC – 3’). The sequencing results were uploaded to Synthego for ICE analysis according to the site instructions^70^.

### Viral spread

To assess the spread of GFP-expressing virus over time, MRC5 fibroblasts with the indicated gene modulations were grown to confluence in a 96-well black-walled plate and infected with the denoted virus and MOI. The plates were imaged daily using the Cytation 5 imaging reader (BioTek) with a 2.5X magnification lens and the YFP channel (469/525 nm). GFP-expressing cells were masked and counted using the Gen5 3.11 software. GFP+ cells were used as a readout for infected cells.

### Fluorescent viral titer quantification

MRC5 fibroblasts with the indicated gene modulations were grown in multiple 96-well plates and infected as indicated. At the time points stated, the corresponding 96-well plate was freeze-thawed three times and stored at -80°C until all time points had been harvested. Meanwhile, MRC5s were grown to confluence in a 384-well plate using FluoroBrite DMEM (Gibco, A1896702) supplemented with 10% FBS and 1X GlutaMAX Supplement (Gibco, 35050061). Once all viral time points were harvested, the 96-well plates were thawed at 37°C. Ten µl of each viral supernatant were transferred from the 96-well plate to the 384-well plate using an Assist Plus pipetting robot (Integra-Biosciences) with a 12-channel VOYAGER electronic pipette (Integra-Biosciences). Each sample was subsequently serially diluted twice, for a total of three wells per viral supernatant.

At 48 hours post-infection, GFP-expressing cells were imaged using the Cytation 5 imaging reader (BioTek) with a 2.5X magnification lens and the YFP channel. GFP+ cells were masked and counted using the Gen5 3.11 software. The number of infected cells per well were considered as ‘infectious units’, and the total infectious units/ml (IU/ml) for each sample was calculated for the appropriate dilution. The titers for each dilution were averaged and used as one sample at the corresponding time point.

### Reagents and treatments

Anisomycin (ANS, Medchemexpress #HY-18982) was solubilized to 1 mM in DMSO and stored as 10 µl aliquots at -80°C. Biotin (Sigma-Aldrich, #B4501-100MG) was solubilized in serum-free DMEM to the indicated concentration immediately before use. Human tumor necrosis factor-α (TNFα, Sigma-Aldrich, H8916-10UG) and interferon gamma (IFNγ, Medchemexpress #HY-P70610-50UG) were solubilized to 10 ug/ml in sterile water and stored as 10 µl aliquots at -80°C.

To determine the effect of ANS on HCMV infection, cells were infected as described above with AD169 at an MOI of 1. After the 2-hour adsorption period, the viral inoculum was aspirated and replaced with serum-free media containing the appropriate dose of ANS. Cell toxicity was determined via Hoechst (Invitrogen, H3570) nuclei count using a Cytation 5 imaging reader (BioTek) with a 2.5X magnification lens. Fluorescence was measured using the DAPI channel, and each nucleus was counted using the Gen5 3.11 (BioTek) software. Viral spread and fluorescent viral replication quantification were performed as described above.

To determine the effect of TNFα and IFNγ on JUNB KO cells, CRISPR EV and JUNB KO cells were treated with 10 ng/ml TNFα or IFNγ for 24 hours.

### BioID sample preparation and pulldown

Samples were treated as described previously^22^. Briefly, MRC5 fibroblasts were mock infected or infected with TurboID-N-UL26wt at an MOI of 3 as described above. At 23 hours post-infection, the medium was replaced with serum-free media containing 1 µM biotin. After a 1-hour incubation, the medium was removed, and the cells were washed twice with cold PBS. Cells were harvested in lysis buffer and clarified. Washed streptavidin-bound magnetic beads (Dynabeads MyOne Streptavidin T1, Invitrogen, 65601) were added to the lysates, and the samples were incubated at room temperature for 1 hour. The samples were placed on a magnetic stand, and the supernatant was removed. The magnetic beads were washed extensively. Protein was eluted from the beads with disruption buffer. The samples were boiled for 5 minutes and centrifuged at 16,000xg for 10 minutes. The supernatant was transferred to a new tube and western blotted as described above.

### Viral DNA accumulation

MRC5 fibroblasts were grown to confluence in 12-well tissue culture plates and infected as described above. At the indicated time points, cells were harvested using 250 µl of QuickExtract DNA extraction solution per well, and samples were processed according to the manufacturer’s instructions. DNA samples were diluted 1:100 in sterile, nuclease-free water and analyzed via RT-qPCR. The relative quantities of IE1 compared to GAPDH were calculated using the 2^-ΔΔCt^ method and normalized to the relative abundance of the CRISPR EV 24hpi AD169 samples. The primers used were: IE1 F (5’ – CCATGTCCACTCGAACCTTAAT – 3’), IE1 R (5’ – TGAACAAGTGACCGAGGATTG – 3’), GAPDH F (5’ – GTCTCCTCTGACTTCAACAGCG – 3’) and GAPDH R (5’ – ACCACCCTGTTGCTGTAGCCAA – 3’).

### Immunofluorescence and Analysis

Glass cover slips (0.2 x 0.2 mm, VWR #48366-227) were sterilized by autoclaving and placed into each well of a 6-well plate. MRC5 fibroblasts with the indicated gene modulation were seeded over the cover slips and allowed to grow to confluence. The cells were infected as described above with the virus and MOI indicated. At 72 hours post-infection, the medium was aspirated from the wells, and the cells were washed 3x with PBS. Fixing solution (2% formaldehyde in PBS) was added to each well and incubated in a fume hood for 15 minutes. The solution was removed, and the cells were washed 3x with PBS. Permeabilization solution (0.1% Triton X-100 (Thermo Scientific Chemicals, A16046-AE), 0.1% SDS in PBS) was added to each well and incubated at room temperature for 15 minutes. The solution was removed, and the cells were washed 3x with IF wash buffer (0.01% Tween-20 (VWR, 97062-332) in PBS). Blocking buffer (2% BSA, 0.05% Tween-20, 0.3% Triton X-100, 5% human serum in PBS) was added to the cells and incubated at 4°C overnight with gentle agitation. The coverslips were transferred to a humidity chamber and placed cell side down onto JUNB primary antibody (Cell Signaling, 3753S) diluted 1:500 in 0.01% Tween-20 in PBS and incubated for 1 hour at room temperature. The coverslips were transferred to a clean 6-well plate and washed 3x with IF wash buffer. The coverslips were transferred to a humidity chamber and placed cell side down onto Alexa Fluor 594 goat anti-rabbit IgG (H+L) secondary antibody (Invitrogen, A-11012) diluted 1:1000 in 0.01% Tween-20 in PBS and incubated covered for 1 hour at room temperature. Coverslips were transferred to a clean 6-well plate and washed 3x with IF wash buffer, with the final wash supplemented with 1:1000 Hoechst 33342 stain. The coverslips were mounted onto glass slides with 90% glycerol and sealed using clear nail polish. The slides were imaged using a Cytation 5 imaging reader (BioTek) with a 40X magnification lens. The JUNB fluorescence was measured using a Texas Red channel with an excitation wavelength of 586 nm and emission wavelength of 647 nm, and the Hoechst fluorescence was measured using a DAPI channel.

For the immunofluorescence visualizing UL44 and JUNB, the cells and coverslips were processed and imaged as above except they were co-incubated with both JUNB primary antibody and UL44 primary antibody (SeraCare, VS-P1202-2). The coverslips were incubated with both Alexa Fluor 594 goat anti-rabbit IgG (H+L) secondary antibody (Invitrogen, A-11012) diluted 1:1000 and Goat anti-Mouse IgG (H+L) Cross-Adsorbed Secondary Antibody, Alexa Fluor™ 405 (Invitrogen, A-31553) diluted 1:400. During the final wash steps, no Hoechst stain was added. UL44 fluorescence was measured using a DAPI channel.

The Gen5 3.11 software was used to calculate the area of JUNB condensates and viral replication compartments. To quantify JUNB condensate area, a primary mask identifying nuclei using the DAPI channel was created. Under this primary mask, a secondary mask identifying JUNB condensates using the Texas Red channel was made. The secondary mask was optimized to include solely the individual puncta of high intensity and outliers were removed. To quantify the area of viral replication compartments, a primary mask identifying UL44 using the DAPI channel was created.

CellProfiler^71^ 4.2.8 was used to calculate the ranked weighted correlation of JUNB and UL44. Briefly, the raw unmerged images were exported from the Gen5 3.11 program and imported into CellProfiler. Images were assigned as either UL44 or JUNB based on their channel. Each image was corrected using the ‘CorrectIlluminationCalculate’ and ‘CorrectIlluminationApply’ modules. Each UL44 object was identified using the corrected images and the ‘IdentifyPrimaryObjects’ module. The Rank Weighted Colocation coefficients between JUNB and UL44 were then calculated using the ‘MeasureColocalization’ module.

O-propargyl-puromycin (OPP) (Click-&-Go^®^ Plus 488 OPP, Vector Laboratories, CCT-1493) was used to measure anisomycin’s effect on protein translation. MRC5 cells were treated with DMSO, 75 nM ANS, 35 µM cycloheximide (CHX), or 750 nM ANS for 24 hours, treated with OPP reagent for 30 minutes, and processed as per the manufacturer’s instructions. Cells were imaged using a Cytation 5 imaging reader (BioTek) with a 40X magnification lens. The mean OPP fluorescence and cell count based on nuclei staining of each field of view were calculated using the Gen5 3.11 software.

### RT-qPCR

RNA was harvested from cells using TRIzol reagent (Invitrogen, 15596018) and isolated using the Direct-zol RNA miniprep plus kit (Zymo Research, R2072). Purified RNA was used to synthesize cDNA using the qScript cDNA synthesis kit (Quantabio, 95047-100). cDNA samples were analyzed using RT-qPCR, and the relative RNA abundances were calculated using the 2^-ΔΔCt^ method. The primers used were: ISG15 F (5’ – GAGCATCCTGGTGAGGAATAAC -3’), ISG15 R (5’ – CGCTCACTTGCTGCTTCA – 3’), IFIT3 F (5’ – CCTGGAATGCTTACGGCAAGCT – 3’), IFIT3 R (5’ – GAGCATCTGAGAGTCTGCCCAA – 3’), BST2 F (5’ – TCTCCTGCAACAAGAGCTGACC – 3’), BST2 R (5’ – TCTCTGCATCCAGGGAAGCCAT – 3’), GAPDH F, and GAPDH R.

### TCID50 Viral Titer

TCID50 was used to titer coronavirus and adenovirus infections. CRISPR EV and JUNB KO cells were infected as described above. At the desired time point post infection, the virus-containing medium was harvested, frozen on dry ice, and stored at -80°C. The viral samples were thawed in a 37°C water bath, vortexed briefly, and centrifuged at 5,000xg for 1.5 minutes to pellet cell debris. The samples were then serially diluted (1:10) eight times, and 50 µl of each dilution was transferred to each well within a row of a 96-well plate containing sub-confluent MRC5 (n = 4) for coronavirus or A549-ACE2^72^ (n = 3, generous gift from the Dewhurst lab, Cellosaurus CVCL_D4Z5) for adenovirus in 50 µl of growth media. Infected wells were identified at 5 or 12-days post infection for coronavirus and adenovirus, respectively. Titers were calculated using a modified Reed and Muench TCID50 calculator from the Lindenbach Lab^73^.

### Statistical analysis

All statistics were conducted using GraphPad Prism v10.3. Figure 1 panels F and G utilized Mann-Whitney tests. The following figure panels utilized unpaired t tests: figure 2 C, D, F, H-J, M, O-Q; figure 3 B, E, G-H; figure 4 D, F, I; figure 5 B, D-H, J-O; figure S3 C. Figure 4 panel H utilized ordinary one-way Anova.

## Supporting information

CRISPR Screen Library

Next-Generation Sequencing Counts

MAGeCK RRA Output

Primers

Minimal Datasets

## Supporting Information

**S#1 Table. CRISPR Screen Library.** Guide sequences used for the CRISPR screen. Experimental guides were generated using CRISPick by the Broad Institute. Non-targeting guides were generated using CRISPOR.

**S#2 Table. Next-Generation Sequencing Counts.** Raw read counts from the initial sequencing of the library prior to generating lentivirus, and raw read counts from the CRISPR screen.

**S#3 Table. MAGeCK RRA Output.** Output from MAGeCK RRA comparing various conditions against t0. Input used both replicates of each condition listed.

**S#4 Table. Primers.** Primers and oligos used for cloning, NGS, sequencing, RNP KO, and RT-qPCR.

**S#5 Data. Minimal datasets.** Raw data used to produce figures for this manuscript. Each sheet is labeled with the corresponding figure panel to which the data pertains to.

## Acknowledgements

We thank The University of Rochester Medical Center’s Genomics Research Center for their assistance with next generation sequencing and analysis.

## Author Contributions

Conceptualization: NW, JM

Methodology: NW, JM

Investigation: NW, JC, XS

Visualization: NW, JM

Supervision: JM

Writing—original draft: NW, JM

Writing—review & editing: NW, JM, JC, XS

---

**Supplemental Figure 1.**
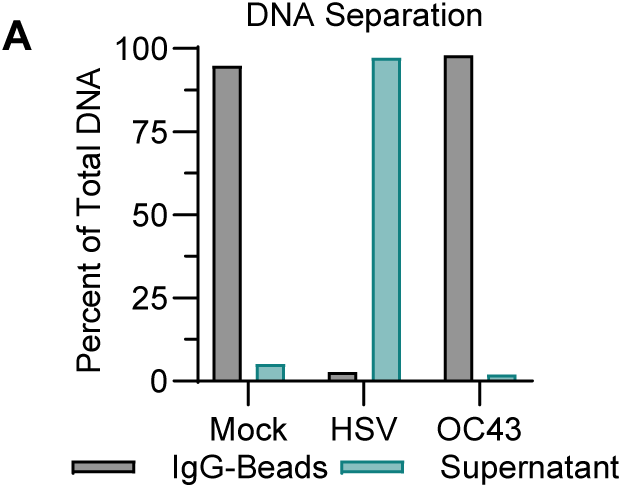
IgG-conjugated bead binding successfully separates HSV-, but not OC43-, infected cells. (A) MRC5 cells were mock-infected or infected with HSV (MOI 5) or OC43 (MOI 5). At 18 hours post infection, cells were harvested using the bead-binding assay as described.

**Supplemental Figure 2.**
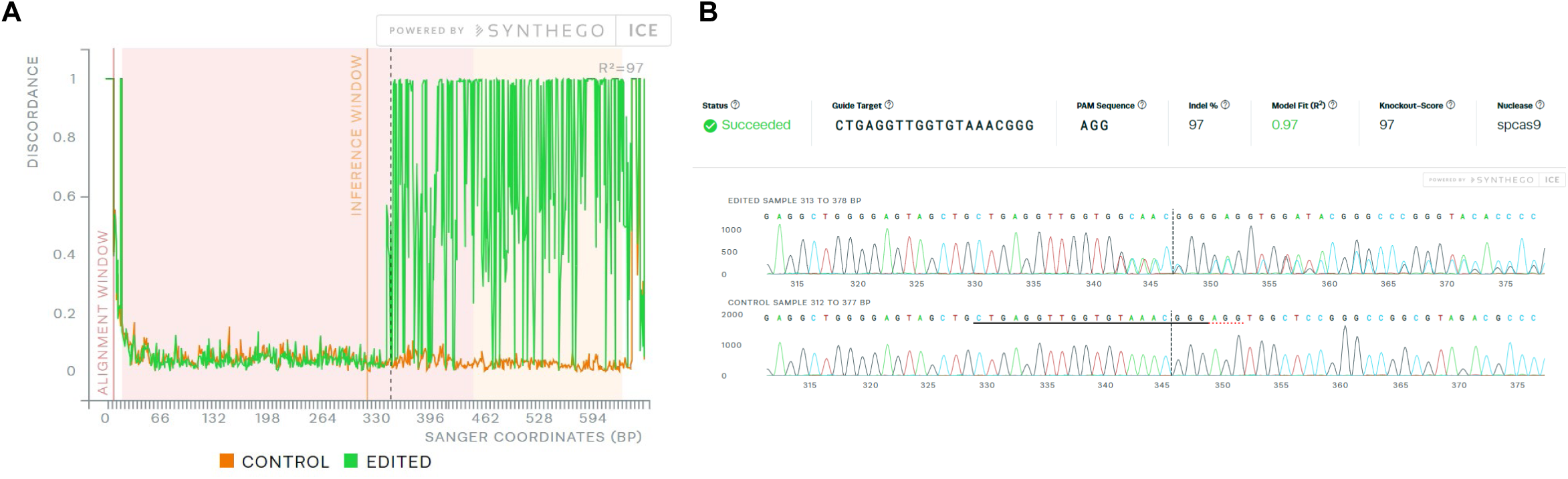
JUNB KO ICE Analysis. (A) Synthego ICE analysis of JUNB KO cells compared to control cells. (B) Sanger sequencing traces of the data presented in (A).

**Supplemental Figure 3.**
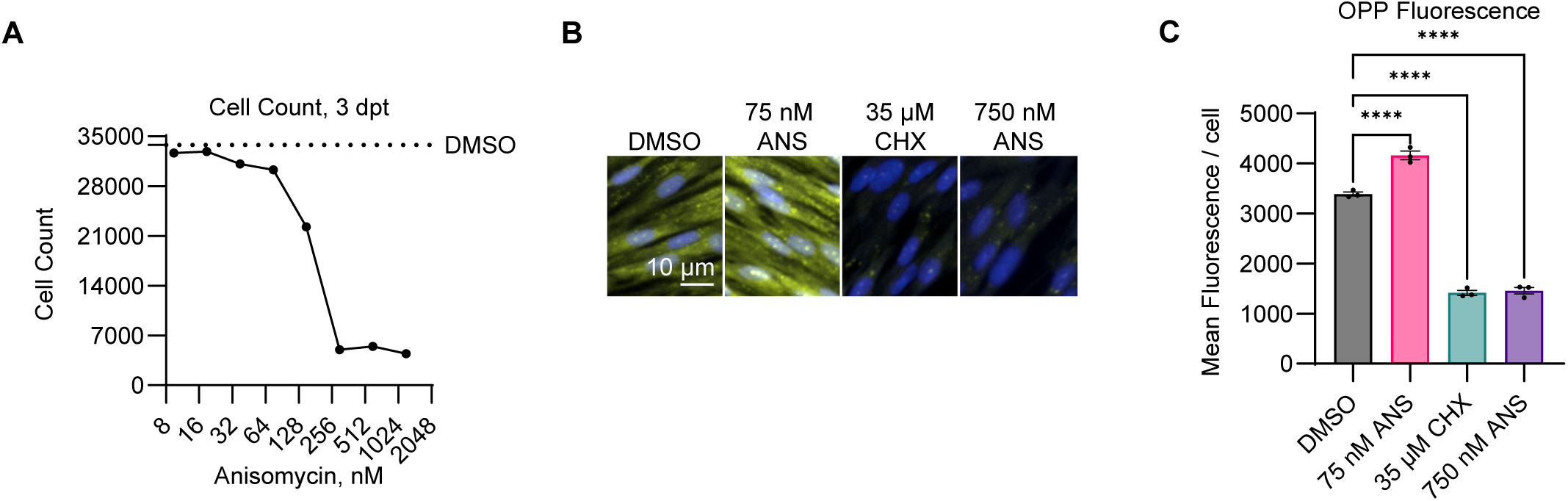
Impact of Anisomycin on protein translation and cell viability. (A) MRC5 cells were treated with Anisomycin (ANS) and stained with Hoechst at 3 days post treatment. Nuclei were counted. (B) MRC5 cells were treated with DMSO, 75 nM ANS, 35 µM cycloheximide (CHX), or 750 nM ANS for 24 hours then treated with OPP reagent for 30 minutes. Cells were stained for OPP (green) and nuclei (Hoechst, blue) and imaged. Brightness and contrast adjusted for visual clarity. (C) Mean OPP fluorescence per cell across three fields of view as in (B). One-way Anova. **** = p<0.0001.

## References

1 Sanson, K. R. et al. Optimized libraries for CRISPR-Cas9 genetic screens with multiple modalities. Nature Communications 9, 5416, doi:10.1038/s41467-018-07901-8 (2018).

2 Doench, J. G. et al. Optimized sgRNA design to maximize activity and minimize off-target effects of CRISPR-Cas9. Nature Biotechnology 34, 184–191, doi:10.1038/nbt.3437 (2016).

3 Puschnik, A. S., Majzoub, K., Ooi, Y. S. & Carette, J. E. A CRISPR toolbox to study virus-host interactions. Nat Rev Microbiol 15, 351–364, doi:10.1038/nrmicro.2017.29 (2017).

4 E, X. & Kowalik, T. F. A Generally Applicable CRISPR/Cas9 Screening Technique to Identify Host Genes Required for Virus Infection as Applied to Human Cytomegalovirus (HCMV) Infection of Epithelial Cells. Methods Mol Biol 2244, 247–264, doi:10.1007/978-1-0716-1111-1_13 (2021).

5 Finkel, Y. et al. A virally encoded high-resolution screen of cytomegalovirus dependencies. Nature, doi:10.1038/s41586-024-07503-z (2024).

6 King, C. R. & Mehle, A. The later stages of viral infection: An undiscovered country of host dependency factors. PLoS Pathog. 16, e1008777, doi:10.1371/journal.ppat.1008777 (2020).

7 Hein, M. Y. & Weissman, J. S. Functional single-cell genomics of human cytomegalovirus infection. Nat. Biotechnol. 40, 391–401, doi:10.1038/s41587-021-01059-3 (2022).

8 Wu, K., Oberstein, A., Wang, W. & Shenk, T. Role of PDGF receptor-α during human cytomegalovirus entry into fibroblasts. Proc Natl Acad Sci U S A 115, E9889–e9898, doi:10.1073/pnas.1806305115 (2018).

9 E, X., et al. OR14I1 is a receptor for the human cytomegalovirus pentameric complex and defines viral epithelial cell tropism. Proc Natl Acad Sci U S A 116, 7043–7052, doi:10.1073/pnas.1814850116 (2019).

10 Cannon, M. J., Schmid, D. S. & Hyde, T. B. Review of cytomegalovirus seroprevalence and demographic characteristics associated with infection. Rev. Med. Virol. 20, 202–213, doi:10.1002/rmv.655 (2010).

11 Zuhair, M. et al. Estimation of the worldwide seroprevalence of cytomegalovirus: A systematic review and meta-analysis. Rev. Med. Virol. 29, e2034, doi:10.1002/rmv.2034 (2019).

12 Fowler, K. B. et al. The outcome of congenital cytomegalovirus infection in relation to maternal antibody status. N. Engl. J. Med. 326, 663–667, doi:10.1056/NEJM199203053261003 (1992).

13 Grgic, I. & Gorenec, L. Human Cytomegalovirus (HCMV) Genetic Diversity, Drug Resistance Testing and Prevalence of the Resistance Mutations: A Literature Review. Trop Med Infect Dis 9, doi:10.3390/tropicalmed9020049 (2024).

14 Limaye, A. P., Babu, T. M. & Boeckh, M. Progress and Challenges in the Prevention, Diagnosis, and Management of Cytomegalovirus Infection in Transplantation. Clin. Microbiol. Rev. 34, doi:10.1128/CMR.00043-19 (2020).

15 Felicia Goodrum, W. B. E. S. M. in Fields Virology: DNA Viruses 389–444.

16 Stamminger, T. et al. Open reading frame UL26 of human cytomegalovirus encodes a novel tegument protein that contains a strong transcriptional activation domain. J. Virol. 76, 4836–4847, doi:10.1128/jvi.76.10.4836-4847.2002 (2002).

17 Lorz, K. et al. Deletion of open reading frame UL26 from the human cytomegalovirus genome results in reduced viral growth, which involves impaired stability of viral particles. J. Virol. 80, 5423–5434, doi:10.1128/JVI.02585-05 (2006).

18 Munger, J., Yu, D. & Shenk, T. UL26-deficient human cytomegalovirus produces virions with hypophosphorylated pp28 tegument protein that is unstable within newly infected cells. J. Virol. 80, 3541–3548, doi:10.1128/JVI.80.7.3541-3548.2006 (2006).

19 Mathers, C., Spencer, C. M. & Munger, J. Distinct domains within the human cytomegalovirus U(L)26 protein are important for wildtype viral replication and virion stability. PLoS One 9, e88101, doi:10.1371/journal.pone.0088101 (2014).

20 Mathers, C., Schafer, X., Martinez-Sobrido, L. & Munger, J. The human cytomegalovirus UL26 protein antagonizes NF-κB activation. J. Virol. 88, 14289–14300, doi:10.1128/JVI.02552-14 (2014).

21 Goodwin, C. M., Schafer, X. & Munger, J. UL26 Attenuates IKKβ-Mediated Induction of Interferon-Stimulated Gene (ISG) Expression and Enhanced Protein ISGylation during Human Cytomegalovirus Infection. J. Virol. 93, doi:10.1128/JVI.01052-19 (2019).

22 Ciesla, J., Huang, K.-L., Wagner, E. J. & Munger, J. A UL26-PIAS1 complex antagonizes anti-viral gene expression during Human Cytomegalovirus infection. PLoS Pathog. 20, e1012058, doi:10.1371/journal.ppat.1012058 (2024).

23 Ren, X. et al. JunB condensation attenuates vascular endothelial damage under hyperglycemic condition. J. Mol. Cell Biol., doi:10.1093/jmcb/mjad072 (2023).

24 Weekes, M. P. et al. Quantitative temporal viromics: an approach to investigate host-pathogen interaction. Cell 157, 1460–1472, doi:10.1016/j.cell.2014.04.028 (2014).

25 Lachmann, A. et al. ChEA: transcription factor regulation inferred from integrating genome-wide ChIP-X experiments. Bioinformatics 26, 2438–2444, doi:10.1093/bioinformatics/btq466 (2010).

26 Li, W. et al. MAGeCK enables robust identification of essential genes from genome-scale CRISPR/Cas9 knockout screens. Genome Biol. 15, 554, doi:10.1186/s13059-014-0554-4 (2014).

27 Terhune, S. et al. Human cytomegalovirus UL38 protein blocks apoptosis. J. Virol. 81, 3109–3123, doi:10.1128/JVI.02124-06 (2007).

28 Sartini, B. L., Wang, H., Wang, W., Millette, C. F. & Kilpatrick, D. L. Pre-Messenger RNA Cleavage Factor I (CFIm): Potential Role in Alternative Polyadenylation During Spermatogenesis1. Biology of Reproduction 78, 472–482, doi:10.1095/biolreprod.107.064774 (2008).

29 Figueroa, J. D. & Hayman, M. J. Differential effects of the Ski-interacting protein (SKIP) on differentiation induced by transforming growth factor-β1 and bone morphogenetic protein-2 in C2C12 cells. Experimental Cell Research 296, 163–172, doi:10.1016/j.yexcr.2004.01.025 (2004).

30 Tu, Y.-T. & Barrientos, A. The Human Mitochondrial DEAD-Box Protein DDX28 Resides in RNA Granules and Functions in Mitoribosome Assembly. Cell Rep. 10, 854–864, doi:10.1016/j.celrep.2015.01.033 (2015).

31 Myers, L. C. & Kornberg, R. D. Mediator of transcriptional regulation. Annu Rev Biochem 69, 729–749, doi:10.1146/annurev.biochem.69.1.729 (2000).

32 Chiu, R., Angel, P. & Karin, M. Jun-B differs in its biological properties from, and is a negative regulator of, c-Jun. Cell 59, 979–986, doi:10.1016/0092-8674(89)90754-x (1989).

33 Eferl, R. & Wagner, E. F. AP-1: a double-edged sword in tumorigenesis. Nat. Rev. Cancer 3, 859–868, doi:10.1038/nrc1209 (2003).

34 Schmidt, D. et al. Critical role for NF-kappaB-induced JunB in VEGF regulation and tumor angiogenesis. EMBO J. 26, 710–719, doi:10.1038/sj.emboj.7601539 (2007).

35 Ren, F.-J., Cai, X.-Y., Yao, Y. & Fang, G.-Y. JunB: a paradigm for Jun family in immune response and cancer. Front. Cell. Infect. Microbiol. 13, 1222265, doi:10.3389/fcimb.2023.1222265 (2023).

36 Hori, T. et al. Molecular mechanism of apoptosis and gene expressions in human lymphoma U937 cells treated with anisomycin. Chemico-Biological Interactions 172, 125–140, doi:10.1016/j.cbi.2007.12.003 (2008).

37 Hazzalin, C. A., Le Panse, R., Cano, E. & Mahadevan, L. C. Anisomycin selectively desensitizes signalling components involved in stress kinase activation and fos and jun induction. Mol. Cell. Biol. 18, 1844–1854, doi:10.1128/mcb.18.4.1844 (1998).

38 Mahadevan, L. C. & Edwards, D. R. Signalling and superinduction. Nature 349, 747–748, doi:10.1038/349747c0 (1991).

39 Ahn, J. H., Jang, W. J. & Hayward, G. S. The human cytomegalovirus IE2 and UL112-113 proteins accumulate in viral DNA replication compartments that initiate from the periphery of promyelocytic leukemia protein-associated nuclear bodies (PODs or ND10). J. Virol. 73, 10458–10471, doi:10.1128/JVI.73.12.10458-10471.1999 (1999).

40 Mariamé, B. et al. Real-time visualization and quantification of human Cytomegalovirus replication in living cells using the ANCHOR DNA labeling technology. J. Virol. 92, JVI.00571-00518, doi:10.1128/JVI.00571-18 (2018).

41 Strang, B. L., Boulant, S., Kirchhausen, T. & Coen, D. M. Host cell nucleolin is required to maintain the architecture of human Cytomegalovirus replication compartments. mBio 3, doi:10.1128/mbio.00301-11 (2012).

42 Liska, O. et al. TFLink: an integrated gateway to access transcription factor-target gene interactions for multiple species. Database (Oxford) 2022, doi:10.1093/database/baac083 (2022).

43 Bianco, C. & Mohr, I. Restriction of Human Cytomegalovirus Replication by ISG15, a Host Effector Regulated by cGAS-STING Double-Stranded-DNA Sensing. Journal of Virology 91, 10.1128/jvi.02483-02416, doi:10.1128/jvi.02483-16 (2017).

44 van den Pol, A. N., et al. Cytomegalovirus induces interferon-stimulated gene expression and is attenuated by interferon in the developing brain. J Virol 81, 332–348, doi:10.1128/jvi.01592-06 (2007).

45 Ashley, C. L., Abendroth, A., McSharry, B. P. & Slobedman, B. Interferon-Independent Upregulation of Interferon-Stimulated Genes during Human Cytomegalovirus Infection is Dependent on IRF3 Expression. Viruses 11, doi:10.3390/v11030246 (2019).

46 Viswanathan, K. et al. BST2/Tetherin enhances entry of human cytomegalovirus. PLoS Pathog 7, e1002332, doi:10.1371/journal.ppat.1002332 (2011).

47 Thomsen, M. K. et al. JUNB/AP-1 controls IFN-gamma during inflammatory liver disease. J Clin Invest 123, 5258–5268, doi:10.1172/JCI70405 (2013).

48 van Dam, H. & Castellazzi, M. Distinct roles of Jun : Fos and Jun : ATF dimers in oncogenesis. Oncogene 20, 2453–2464, doi:10.1038/sj.onc.1204239 (2001).

49 Schraml, B. U. et al. The AP-1 transcription factor Bati controls T(H)17 differentiation. Nature 460, 405–409, doi:10.1038/nature08114 (2009).

50 Eferl, R. & Wagner, E. F. AP-1: a double-edged sword in tumorigenesis. Nature Reviews Cancer 3, 859–868, doi:10.1038/nrc1209 (2003).

51 Ishida, T., Nakajima, T., Kudo, A. & Kawakami, A. Phosphorylation of Junb family proteins by the Jun N-terminal kinase supports tissue regeneration in zebrafish. Dev Biol 340, 468–479, doi:10.1016/j.ydbio.2010.01.036 (2010).

52 Yamaguchi, N. et al. c-Abl-mediated tyrosine phosphorylation of JunB is required for Adriamycin-induced expression of p21. Biochem J 471, 67–77, doi:10.1042/bj20150372 (2015).

53 Garaude, J. et al. SUMOylation regulates the transcriptional activity of JunB in T lymphocytes. J Immunol 180, 5983–5990, doi:10.4049/jimmunol.180.9.5983 (2008).

54 Ungureanu, D. et al. PIAS proteins promote SUMO-1 conjugation to STAT1. Blood 102, 3311–3313, doi:10.1182/blood-2002-12-3816 (2003).

55 Liu, B. et al. PIAS1 selectively inhibits interferon-inducible genes and is important in innate immunity. Nature immunology 5, 891–898, doi:10.1038/ni1104 (2004).

56 Ciesla, J., Huang, K. L., Wagner, E. J. & Munger, J. A UL26-PIAS1 complex antagonizes anti-viral gene expression during Human Cytomegalovirus infection. PLoS Pathog 20, e1012058, doi:10.1371/journal.ppat.1012058 (2024).

57 Yu, D., Silva, M. C. & Shenk, T. Functional map of human cytomegalovirus AD169 defined by global mutational analysis. Proc. Natl. Acad. Sci. U. S. A. 100, 12396–12401, doi:10.1073/pnas.1635160100 (2003).

58 Etienne, L., et al. Visualization of herpes simplex virus type 1 virions using fluorescent colors. J. Virol. Methods 241, 46–51, doi:10.1016/j.jviromet.2016.12.012 (2017).

59 Rodríguez-Sánchez, I., Schafer, X. L., Monaghan, M. & Munger, J. The Human Cytomegalovirus UL38 protein drives mTOR-independent metabolic flux reprogramming by inhibiting TSC2. PLOS Pathogens 15, e1007569, doi:10.1371/journal.ppat.1007569 (2019).

60 Zhu, H., Shen, Y. & Shenk, T. Human cytomegalovirus IE1 and IE2 proteins block apoptosis. J Virol 69, 7960–7970, doi:10.1128/jvi.69.12.7960-7970.1995 (1995).

61 Silva, M. C., Yu, Q. C., Enquist, L. & Shenk, T. Human cytomegalovirus UL99-encoded pp28 is required for the cytoplasmic envelopment of tegument-associated capsids. J Virol 77, 10594–10605, doi:10.1128/jvi.77.19.10594-10605.2003 (2003).

62 Nobre, L. V. et al. Human cytomegalovirus interactome analysis identifies degradation hubs, domain associations and viral protein functions. Elife 8, doi:10.7554/eLife.49894 (2019).

63 Kuleshov, M. V. et al. Enrichr: a comprehensive gene set enrichment analysis web server 2016 update. Nucleic Acids Res. 44, W90–97, doi:10.1093/nar/gkw377 (2016).

64 Concordet, J.-P. & Haeussler, M. CRISPOR: intuitive guide selection for CRISPR/Cas9 genome editing experiments and screens. Nucleic Acids Research 46, W242–W245, doi:10.1093/nar/gky354 (2018).

65 Kechin, A., Boyarskikh, U., Kel, A. & Filipenko, M. cutPrimers: A New Tool for Accurate Cutting of Primers from Reads of Targeted Next Generation Sequencing. J Comput Biol 24, 1138–1143, doi:10.1089/cmb.2017.0096 (2017).

66 Langmead, B., Trapnell, C., Pop, M. & Salzberg, S. L. Ultrafast and memory-efficient alignment of short DNA sequences to the human genome. Genome Biology 10, R25, doi:10.1186/gb-2009-10-3-r25 (2009).

67 Liao, Y., Smyth, G. K. & Shi, W. The R package Rsubread is easier, faster, cheaper and better for alignment and quantification of RNA sequencing reads. Nucleic Acids Res 47, e47, doi:10.1093/nar/gkz114 (2019).

68 Ciesla, J., Moreno, I., Jr. & Munger, J. TNFα-induced metabolic reprogramming drives an intrinsic anti-viral state. PLoS Pathog. 18, e1010722, doi:10.1371/journal.ppat.1010722 (2022).

69 Horlbeck, M. A. et al. Compact and highly active next-generation libraries for CRISPR-mediated gene repression and activation. Elife 5, doi:10.7554/eLife.19760 (2016).

70 Conant, D. et al. Inference of CRISPR Edits from Sanger Trace Data. Crispr j 5, 123–130, doi:10.1089/crispr.2021.0113 (2022).

71 Stirling, D. R., et al. CellProfiler 4: improvements in speed, utility and usability. BMC Bioinformatics 22, 433, doi:10.1186/s12859-021-04344-9 (2021).

72 Chang, C.-W. et al. A newly engineered A549 cell line expressing ACE2 and TMPRSS2 is highly permissive to SARS-CoV-2, including the delta and omicron variants. Viruses 14, 1369, doi:10.3390/v14071369 (2022).

73 Lindenbach, B. D. Measuring HCV infectivity produced in cell culture and in vivo. Methods Mol. Biol. 510, 329–336, doi:10.1007/978-1-59745-394-3_24 (2009).

